# Rare copy number variants in *NRXN1* and *CNTN6* increase risk for Tourette syndrome

**DOI:** 10.1101/062471

**Authors:** Alden Y. Huang, Dongmei Yu, Lea K. Davis, Jae-Hoon Sul, Fotis Tsetsos, Vasily Ramensky, Ivette Zelaya, Eliana Marisa Ramos, Lisa Osiecki, Jason A. Chen, Lauren M. McGrath, Cornelia Illmann, Paul Sandor, Cathy L. Barr, Marco Grados, Harvey S. Singer, Markus M. Noethen, Johannes Hebebrand, Robert A. King, Yves Dion, Guy Rouleau, Cathy L. Budman, Christel Depienne, Yulia Worbe, Andreas Hartmann, Kirsten R. Muller-Vahl, Manfred Stuhrmann, Harald Aschauer, Mara Stamenkovic, Monika Schloegelhofer, Anastasios Konstantinidis, Gholson J. Lyon, William M. McMahon, Csaba Barta, Zsanett Tarnok, Peter Nagy, James R. Batterson, Renata Rizzo, Danielle C. Cath, Tomasz Wolanczyk, Cheston Berlin, Irene A. Malaty, Michael S. Okun, Douglas W. Woods, Elliott Rees, Carlos N. Pato, Michele T. Pato, James A Knowles, Danielle Posthuma, David L. Pauls, Nancy J. Cox, Benjamin M. Neale, Nelson B. Freimer, Peristera Paschou, Carol A. Mathews, Jeremiah M. Scharf, Giovanni Coppola

## Abstract

Tourette syndrome (TS) is highly heritable, although identification of its underlying genetic cause(s) has remained elusive. We examined a European ancestry sample composed of 2,435 TS cases and 4,100 controls for copy-number variants (CNVs) using SNP microarrays and identified two genome-wide significant loci that confer a substantial increase in risk for TS (*NRXN1*, OR=20.3, 95%CI [2.6-156.2], p=6.0 × 10^−6^; *CNTN6*, OR=10.1, 95% CI [2.3-45.4], p=3.7 × 10^−5^). Approximately 1% of TS cases carried one of these CNVs, indicating that rare structural variation contributes significantly to the genetic architecture of TS.

---

Tourette syndrome (TS) is a complex developmental neuropsychiatric disorder of childhood onset characterized by multiple motor and vocal tics, with an estimated population prevalence of 0.3-0.9%^1^. Twin and family-based studies of TS have repeatedly demonstrated that it is highly heritable (e.g., h^2^ of 0.77 in a recent analysis of the Swedish National Patient Register^2^), while analysis of genome-wide SNP data suggests that TS risk is highly polygenic and distributed across both common and rare variation^3^. To date, the TS samples with available genome-wide genotyping have been inadequately sized for common variant association studies of a complex trait. To further characterize the genetic influences on TS, we assessed the impact of rare CNVs on disease risk, as it has been shown repeatedly that such variants contribute to susceptibility for other heritable neurodevelopmental disorders, including intellectual disability (ID), autism spectrum disorder (ASD) and schizophrenia (SCZ)^4^.

We genotyped TS cases and ancestry-matched controls on the same genome-wide SNP array platform (Illumina OmniExpress, Supplementary Table 1a). Following standard quality control (QC) steps (Supplementary Table 1b, Supplementary Figure 1, and Supplementary Text), including genotype-based determination of ancestry (Supplementary Figure 2), we analyzed CNVs in a SNP dataset of 6,535 unrelated European ancestry samples, including 2,435 individuals diagnosed with TS (by DSM-IV-TR criteria) and 4,100 healthy controls. To improve specificity, we generated CNV calls with two widely-used Hidden Markov Model (HMM)-based segmentation algorithms, PennCNV^5^ and QuantiSNP^6^ and retained the intersection of CNVs detected by both methods. We also conducted an additional QC step to test for any differential sensitivity in CNV detection between cases and controls, both within and across batches and sites, by analyzing CNV calls from 11 common HapMap3 CNVs using a sensitive locus-specific intensity clustering method, generating a total of 4,758 non-reference CNV calls across all samples (Supplementary Figure 3). Comparison of these genotypes with our consensus HMM-based calls confirmed the absence of any differential bias in the sensitivity of CNV detection between phenotypic groups whether assessed across all loci (p=0.53, Fisher’s exact test) or between individuals (p=0.15, Welch’s *t*-test, Supplementary Table 4 and Supplementary Text).

In total, we resolved 8,365 rare (as defined by a minor allele frequency [MAF] < 1% across all samples [50% reciprocal overlap]) CNV calls of at least 50 kbps in length and spanning a minimum of 10 probes. We assessed global CNV burden in terms of the number of CNVs, total CNV length, and the number of genes affected by CNVs, stratified by CNV type, size, and frequency. When considering all CNVs, we observed a modest but significant enrichment in TS across all metrics of burden for single-occurrence events (or singletons, corresponding to a MAF of approximately 0.00015) only (OR per CNV of 1.09 [1.01-1.18], p=0.03; OR per 100kb of 1.022 [1.006-1.035], p=6 × 10^−3^; OR per gene of 1.016 [1.004-1.030], p=0.01; Supplementary Table 5). In general, CNV burden for TS was greater with increasing event size and rarity, with the most substantial effect seen among large singleton deletions (>1 Mb), with an OR per CNV of 2.82 [1.36-6.18], p=7 × 10^−3^ (Figure 1).

**Figure 1.**
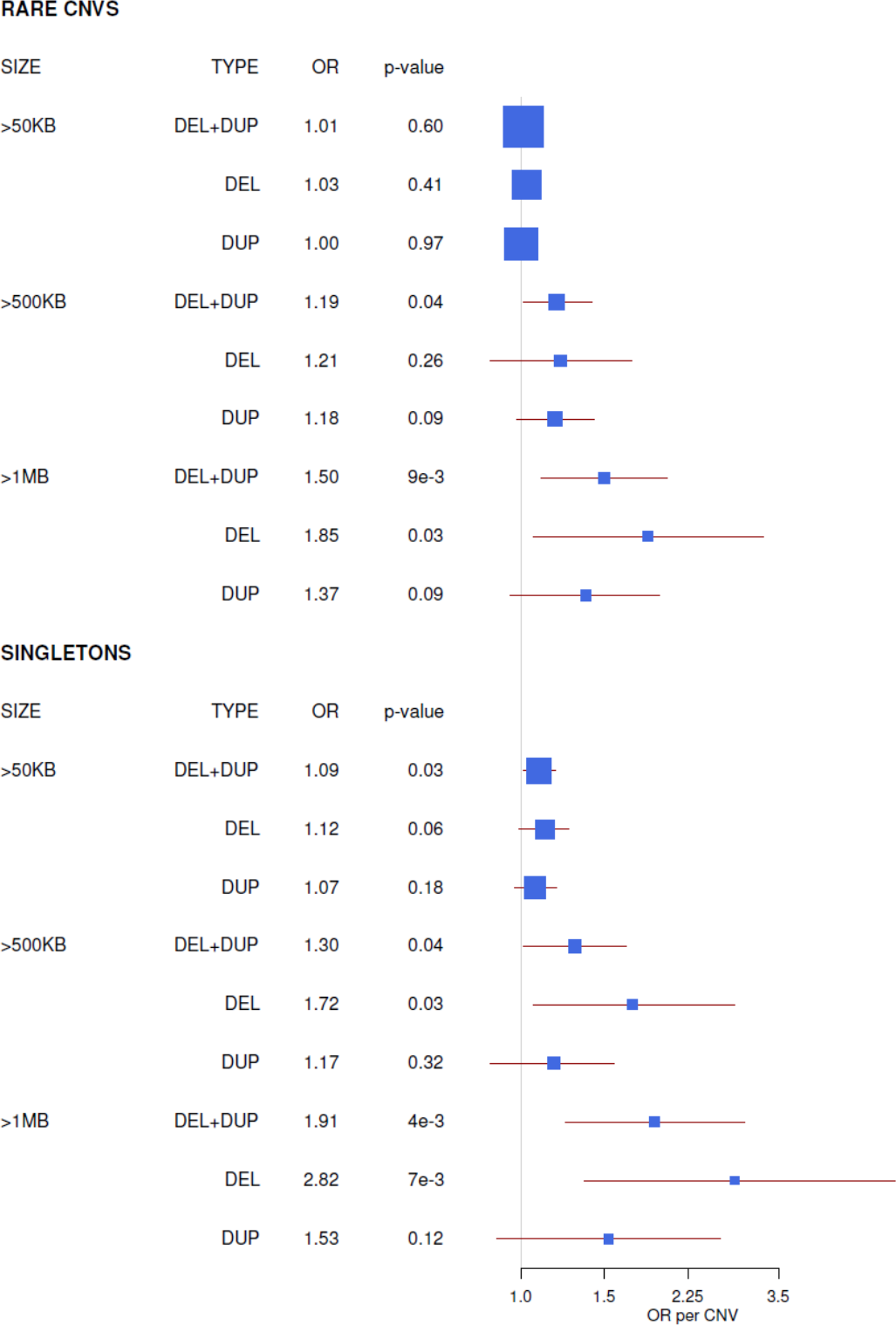
Global CNV burden in Tourette Syndrome (TS) increases with both size and rarity of events. Forest plots show OR and 95% CI estimates for an increased risk for TS per CNV, determined by fitting a logistic regression model of phenotype status against CNV count with significant covariates (Supplementary methods). All CNVs considered are >50kb in length and span a minimum of 10 probes. Rare CNVs denote events with a MAF < 0.01; singletons occur only once in either a case or control, corresponding to a MAF of approximately 0.00015 in this study. P-values are calculated using the likelihood ratio test. Box sizes are proportional to precision.

We next evaluated the dataset for possible enrichment of rare CNVs at specific loci, conducting a point-wise (segmental) test of association, treating deletions and duplications independently. As non-overlapping CNVs that affect the same gene would be unaccounted for by segmental assessments of enrichment, we also performed a complementary test collapsed on individual genes conditioned on exonic CNVs affecting protein-coding genes. In contrast to genome-wide association studies of SNPs, there is no generally accepted threshold to indicate genome-wide significance for CNVs. Therefore, for both tests, we established locus-specific (P_locus_) and genome-wide corrected (P_corr_) p-values empirically through 1,000,000 permutations, using the max(T) method to control for familywise error rate (FWER)^7^. Both tests converged on the same two distinct loci, one for deletions and another for duplications, which were enriched among TS cases and survived correction for multiple testing.

For deletions, the peak segmental association signal was located at rs13418185 (P_locus_=6 × 10^−6^, P_corr_=8.6 × 10^−4^, Figure 2a), corresponding to heterozygous losses across the first three exons of *NRXN_1_*, found exclusively among TS cases (N=10, Figure 2a and b). In the gene-based test of genome-wide exonic CNVs, *NRXN_1_* deletions were the most significant association (Supplementary Table 6 and Supplementary Figure 6), representing 12 cases (0.49%) and a single control (0.02%); OR=20.3 [2.6-156.2]; P_locus_=6.2 × 10^−5^; P_corr_=6.7 × 10^−4^. Consistent with deletions previously identified for this gene in ASD, SCZ, and epilepsy, these exon-spanning CNVs clustered at the 5’ end of the gene and predominantly affected the α isoform of *NRXN_1_^8^*.

**Figure 2.**
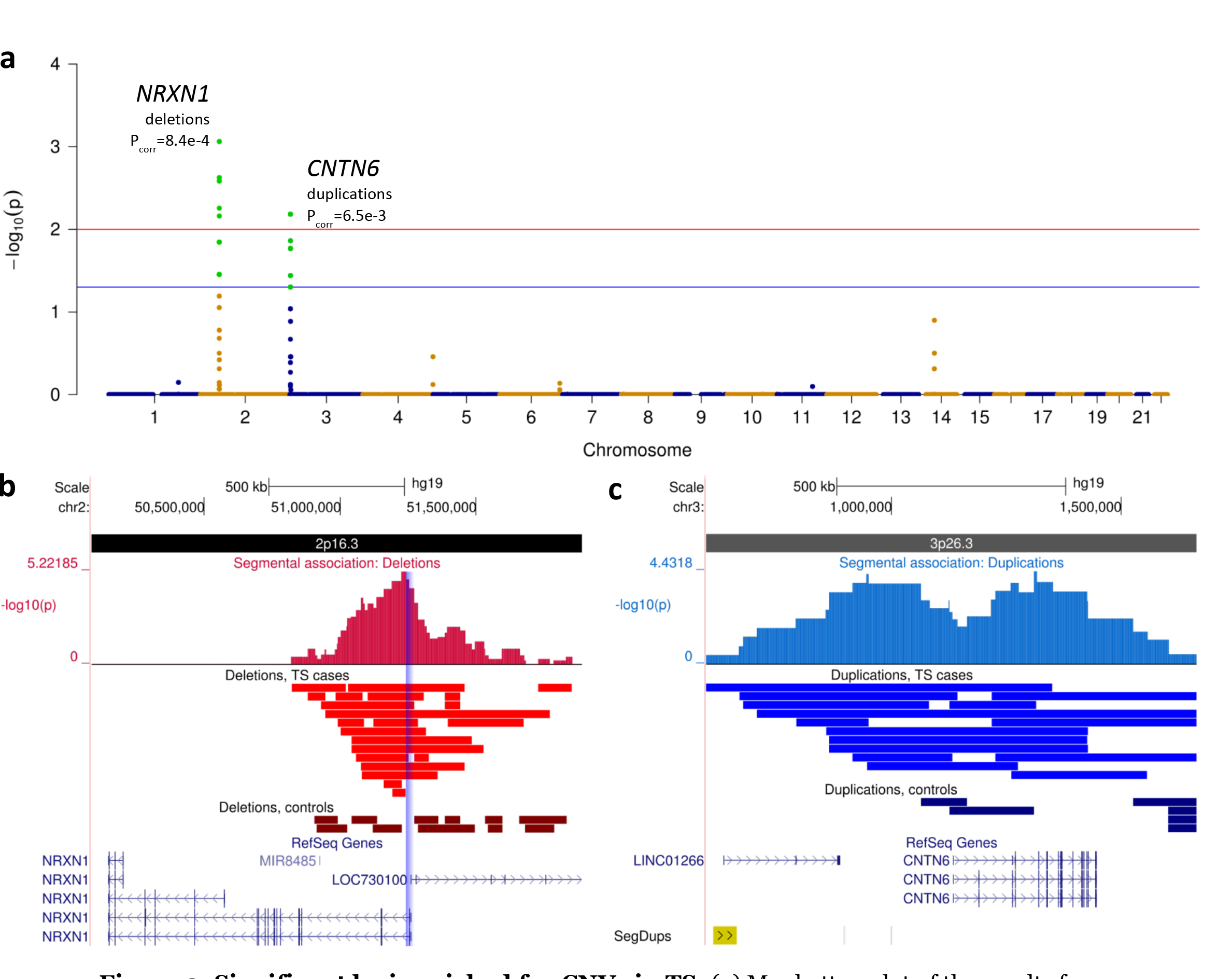
Significant loci enriched for CNVs in TS. (a) Manhattan plot of the results from segmental association tests reveal two genome-wide significant loci: deletions at *NRXN_1_* and duplications at *CNTN6*. Association tests were conducted separately for gains and losses; for clarity, p-values displayed are empirically corrected for FWER genome-wide using the max(T) method with 1,000,000 permutations and plotted together here. Significance levels representing an a of 0.05 and 0.01 are depicted by blue and red lines, respectively. (b) Heterozygous deletions show a peak of segmental association near the 5’ end of *NRXN_1_* (p=6.0 × 10^−6^). Deletions spanning the first three exons are found in 10 cases and no controls (blue-shaded bar). Deletions affecting any exon (Supplementary Figure 6) were found in 12 cases (0.49%) and 1 control (0.03%), corresponding to an OR=20.3, 95% CI (2.6-156.2). (c) Structural variation at the *CNTN6* locus. For duplications, the peak of segmental association was located within the *CNTN6* gene (p=3.7 × 10^−5^). Duplications overlapping this gene correspond to an OR=10.1, 95% CI (2.3-45.4), with heterozygous gains found in 12 cases (0.49%) and 2 controls (0.05%). CNVs overlapping *CNTN6* are considerably larger in cases compared to controls (640.0 vs. 142.9 kb, on average).

The most significant segmental association with a duplication was located within the *CNTN6* gene at rs4085434 (P_locus_=3.7 × 10^−5^, P_corr_=6.5 × 10^−3^) with a secondary peak located directly upstream (P_locus_=5.4 × 10^−5^, P_corr_=6.5 × 10^−3^, Figure 2a and c). Closer inspection of the locus revealed an enrichment of large duplications spanning this gene. A gene-based test determined that duplications overlapping *CNTN6* correspond to an OR=10.1 [2.3-45.4] for TS (P_locus_=2.5 × 10^−4^, P_corr_=8.3 × 10^−3^), with gains found in 12 cases (0.49%) and 2 controls (0.05%) (Supplementary Table 6 and Supplementary Figure 7). All duplications detected across *CNTN6* were heterozygous and spanned exons.

No other loci were significant after controlling for FWER, under either segmental or gene-based tests of association. We obtained similar results after pair-matching each individual case with its closest ancestrally matched control, demonstrating that these results are robust and not the result of inter-European population stratification or case-control sample biases (Supplementary Text and Supplementary Figure 8). Furthermore, we observed no significant enrichment of any CNVs among controls.

Excluding these two genome-wide significant loci, we conducted a secondary analysis testing for an increased burden among 27 rare, recurrent CNVs previously associated with various neurodevelopmental/neuropsychiatric disorders. We observed no nominally significant enrichment, either considering these CNVs individually (Supplementary Table 7) or in concert (P=1.0, 2-sided Fisher’s exact test).

Although previous studies have reported heterozygous exonic *NRXN_1_* deletions in 4 TS patients^9,10^, the small sample sizes in these prior studies precluded any definitive association of this deletion with TS. Here, we demonstrate that exonic deletions affecting *NRXN_1_*, particularly those spanning exons 1-3, confer a substantial increase in risk for the disorder. Of note, among the 12 TS cases with exonic *NRXN_1_* deletions, four had another previously diagnosed neurodevelopmental disorder (NDD), including two with ASD (Supplementary Table 8). The association of *NRXN_1_* deletions with different neurodevelopmental disorders represents one of the most consistent findings regarding CNVs in neuropsychiatry^8,11,12^. Our data suggests an approximately two-fold higher prevalence of exonic *NRXN_1_* deletions in TS compared to other neuropsychiatric disorders^12^, although much larger replication cohorts will be necessary to affirm this apparent comparative enrichment. Despite the diverse clinical presentation exhibited by *NRXN_1_* deletion carriers, *in vitro* models using human neurons differentiated from induced pluripotent stem cells have shown independent lines carrying different exonic deletions in the NRXN_1_-a isoform exhibit markedly similar defects in synaptic transmission^13^. NRXN_1_-α interactions are also critical for thalamocortical synaptogenesis and plasticity^14^, underscoring a potential mechanism for its repeated association to developmental neuropsychiatric disorders.

Like *NRXN_1_*, *CNTN6* encodes a cell-adhesion molecule that has been shown to promote neurite outgrowth. On the basis of structural variation, *CNTN6* has been proposed as a candidate gene for intellectual disability and/or developmental delay^15,16^, and deletions affecting *CNTN6* are significantly enriched in ASD^17^. However, none of the subjects with *CNTN6* duplications identified here had a known NDD (Supplementary Table 8). Notably, the *CNTN6* duplications identified in our sample are considerably larger in TS cases compared to controls (641.0 vs. 142.9 kbp). 9 out of 12 TS carriers harbor a duplication exceeding 500 kbp in length, while both of the *CNTN6* duplications found in controls were less than 200 kbp. Furthermore, in a previous CNV study of 1,086 TS cases and 1,789 controls unrelated to the samples used in the current analysis, duplications directly upstream of *CNTN6* demonstrated the greatest enrichment^18^, reinforcing a possible pathogenic significance of *CNTN6* duplications to this disorder. Consistent with northern blot analysis in the adult human nervous system^19^, examination of human brain RNAseq data from the BrainSpan project indicates that *CNTN6* is widely expressed postnatally with highest expression seen both prenatally and postnatally in the cerebellum and mediodorsal thalamus, with additional focal expression in mid-gestational frontal and sensorimotor cortex (Supplementary Figure 9). The thalamus has long been a region of proposed involvement in TS based on multiple levels of evidence including thalamic lesions, human neurophysiology studies, and by recent treatments successes using deep-brain stimulation^20,21^. The cerebellum has also recently been implicated in TS by functional magnetic resonance imaging^22^.

In summary, we have conducted the largest survey of structural variation in TS to date. We identified two genome-wide significant loci that are enriched for rare CNVs in TS: deletions in *NRXN_1_* and duplications in *CNTN6*. Approximately 1% of TS cases carry a CNV in either gene. Furthermore, we demonstrate a significant increase in global CNV burden, primarily for large, extremely rare deletions. This result suggests that additional CNVs that confer susceptibility to TS remain, but their discovery will likely require substantial increases in sample size.

## Summarized Methods

### Sample ascertainment and data generation

TS cases were ascertained through 21 sites across North America, Europe, and Israel through either specialty clinics or a web-based recruitment effort using a validated diagnostic instrument (TICS Inventory, Supplementary Text). A definite DSM-IV-TR diagnosis of TS, determined by an expert clinician, was a requisite for study inclusion. Unselected control samples were collected in conjunction with TS cases. Additional unscreened controls were obtained from four external studies, and SNP data was generated for all samples on the Illumina OmniExpress platform (Supplementary Text and Supplementary Table 1) according to the manufacturer’s specifications. Raw intensities were obtained using GenomeStudio (Illumina). Quality Control (QC, Supplementary Text and Supplementary Table 1b) was conducted using PLINK v. 1.90, Perl, and R scripts. Samples were further excluded if they were of discrepant or indeterminate genetic sex or were outliers based on heterozygosity (Supplementary Figure 2a). When samples exhibited an excessive amount of cryptic relatedness (PI-HAT > 0.185), only the sample with the higher call rate was retained. In addition, control samples that exhibited an excessive amount of cryptic relatedness to individuals clinically diagnosed with a neuropsychiatric phenotype were also removed.

### Ancestry inference and matching

Genotype data was combined with data from publicly available HapMap samples of European, African, and Asian continental ancestry (Illumina). All available European (EU) population samples from the 1000 Genomes Project were also included to establish an appropriate calibration threshold for EU ancestry designation. A total of 19,024 LD-independent markers were used for ancestry inference, and samples were excluded if they contained > 0.0985 non-EU ancestry as determined using fastStructure (Supplementary Figure 2).

### CNV calling

Only SNP assays common to all versions of the OmniExpress arrays were used for CNV detection (n=689,077) to mitigate any disparity in CNV detection due to probe coverage. Raw CNV calls were generated on all autosomal chromosomes using PennCNV and QuantiSNP. In addition to hard cutoffs used to flag problematic assays, samples were excluded if they represented outliers in a number of CNV quality metrics (determined as mean ±3 SD or by manual inspection, Supplementary Figure 1). Rare CNVs, defined by a prevalence of <1% across all samples, were validated with an alternative locus specific CNV genotyping algorithm that considers normalized, median-summarized intensity values across each putative CNV region. An overview of the CNV processing pipeline is presented in Supplementary Figure 1 and described in detail in the Supplementary Text.

### Burden analysis of global CNV burden

Under a logistic regression model, we assessed for global CNV burden as measured by the total number of CNVs, cumulative CNV length, or number of genes spanned by CNVs, including covariates found to be significantly associated with these burden metrics (Supplementary Text and Supplementary Table 9). Odds ratios indicate an increase in risk for TS per unit of CNV burden. P-values were calculated using the likelihood ratio test.

### Locus-specific tests of association

The segmental test of association was performed at all unique CNV breakpoints. For gene-based association tests, we considered only CNVs spanning exons of coding genes as defined by Refseq annotation. Significance for both tests of association was determined by 1,000,000 permutations of phenotype labels. In each case, both locus-specific and genome-wide corrected p-values were obtained using the max(T) permutation method as implemented in PLINK v1.07, which controls for family-wise error rate by comparing the locus-specific test statistic to all test statistics genome-wide within each permutation.

### Analysis of known neuropsychiatric susceptibility loci

A list of known CNVs with strong evidence of association to various neuropsychiatric disorders, including ASD, ID/DD, SCZ, and BD was assembled from the literature^4^. For recombination hotspots, a CNV was counted if it overlapped with the reported region by at least 50%. Singlegene associated CNVs were considered if they shared overlap to annotated gene boundaries as annotated in RefSeq. Locus-specific P-values were determined by 100,000 permutations of phenotype labels.

## URLs

PennCNV, http://penncnv.openbioinformatics.org; QuantiSNP, https://sites.google.com/site/quantisnp/home; HapMap3, ftp://ftp.ncbi.nlm.nih.gov/hapmap; 1000 Genomes Project; http://www.1000genomes.org; BrainSpan, http://www.brainspan.org

## ACKNOWLEDGMENTS

The authors wish to thank all the patients with Tourette Syndrome and their families, as well as unaffected volunteers, who generously agreed to participate in this study. This study was supported by the US National Institutes of Health U01 NS040024 to Drs. Pauls, Mathews, and Scharf and the Tourette Syndrome Association International Consortium for Genetics, ARRA Grant NS040024-09S1, K23 MH085057, and K02 NS085048 to Dr. Scharf, ARRA Grants NS040024-07S1 and NS016648 to Dr. Pauls, MH096767 to Dr. Mathews and NINDS Informatics Center for Neurogenetics and Neurogenomics grant P30 NS062691 to Drs. Coppola and Freimer, by grants from the Tourette Association of America to Drs. Paschou, Pauls, Mathews, and Scharf and from the German Research Society to Dr. Hebebrand.

### AUTHOR CONTRIBUTIONS

All authors were involved in the conception and design of the genetic study. A.H., P.P., C.A.M., J.M.S., and G.C. designed and oversaw the analyses. A.H., D.Y., L.K.D., J.H.S., F.T., V.R., I.Z., E.M.R., L.O., J.A.C., L.M.M., B.M.N., N.B.F., P.P., C.A.M., J.M.S., and G.C. participated in the conduct of the analyses. Major contributions to writing and editing were made by A.H., C.A.M., J.M.S, and G.C. All authors assisted with critically revising the manuscript.

### COMPETING FINANCIAL INTERESTS

The authors declare competing financial interest: details are available in the online version of the paper.

P.S., M.G., H.S.S., R.A.K., Y.D., G.R., C.L.B., G.L., W.M.M., D.L.P, N.J.C., N.B.F., P.P., C.A.M. and J.M.S have received research funding from the Tourette Association of America (TAA). J.M.S., C.A.M. have received travel support from the TAA and serve on the TAA Scientific Advisory Board. J.M.S. has also received consulting fees from Nuvelation Pharma, Inc. P.S. received unrestricted Educational Grants in support of conferences he organized from Purdue and Shire, a CME speaker fee from Purdue University, industry sponsored clinical trial support from Otsuka and is a member of the Data Safety Monitoring Committee at Psyadon. C.L.B. has received funding for clinical trials from Psyadon Pharmaceuticals, Neurocrine Pharmaceuticals, Synchroneuron Pharmaceuticals, AUSPEX Pharmaceuticals, and TEVA Pharmaceuticals. She was a paid speaker for the TAA CDC program and a paid consultant for Bracket eCOA. I.A.M. has participated in research funded by the National Parkinson Foundation, TAA, Abbvie, Auspex, Biotie, Michael J. Fox Foundation, Neurocrine, Pfizer, and Teva, but has no owner interest in any pharmaceutical company. I.A.M. has been reimbursed for speaking for the National Parkinson Foundation and TAA. M.S.O. serves as a consultant for the National Parkinson Foundation, and has received research grants from NIH, NPF, the Michael J. Fox Foundation, the Parkinson Alliance, Smallwood Foundation, the Bachmann-Strauss Foundation, the TAA, and the UF Foundation. M.S.O’s DBS research is supported by: R01 NR014852 and R01NS096008. He has previously received honoraria, but in the past >60 months has received no support from industry. M.S.O. has received royalties for publications with Demos, Manson, Amazon, Smashwords, Books4Patients, and Cambridge, is an associate editor for New England Journal of Medicine Journal Watch Neurology, has participated in CME and educational activities on movement disorders in the last 36 months sponsored by PeerView, Prime, QuantiaMD, WebMD, Medicus, MedNet, Henry Stewart, and by Vanderbilt University. The institution and not M.S.O. receives grants from Medtronic, Abbvie, Allergan, and ANS/St. Jude, and the PI has no financial interest in these grants. D.W.W. has received royalties from Guilford Press and Oxford University Press, speaking honoraria from the TAA and serves on the TAA Medical Advisory Board. All other authors have no competing financial interests to declare.

## Supplementary Text

### Sample Ascertainment

Tourette Syndrome (TS) cases were ascertained primarily from TS specialty clinics through sites distributed throughout North America, Europe and Israel as part of an ongoing collaborative effort by the Tourette Syndrome Association International Consortium for Genetics (TSAICG) as described in detail elsewhere^23^. Subjects were assessed for a lifetime diagnosis of TS, Obsessive Compulsive Disorder (OCD) and Attention-Deficit/Hyperactivity Disorder (ADHD) using a standardized and validated semi-structured direct interview (TICS Inventory^24^,^25^). An additional 628 cases were collected at 9 TS specialty clinics in Austria, Canada, France, Germany, Greece, Hungary, Italy and the Netherlands by expert clinicians using Tourette Syndrome Study Group criteria for Definite TS (DSM-IV TS diagnosis plus tics observed by a trained clinician) as well as DSM-IV diagnostic criteria for OCD and ADHD, along with 610 ancestry-matched controls as described previously^26^. Formal standardized assessments were not conducted for other neurodevelopmental disorders (NDDs) such as Intellectual Disability/Developmental Delay (ID/DD), Autism Spectrum Disorder (ASD), or schizophrenia/childhood psychosis; however, participants and their parents were asked about the presence of established or suspected NDD diagnoses.

Lastly, in an effort to greatly increase sample size for TS genetic studies in a cost-effective manner, additional TS case samples were obtained through web-based recruitment of individuals with a prior clinical diagnosis of TS who subsequently completed an online questionnaire that we have validated against the gold-standard TS structured diagnostic interview with nearly 100% concordance for all inclusion/exclusion criteria as well as high correct classification rates for DSM-IV diagnoses of OCD and ADHD^25^,^27^. Individuals for web-based screening were solicited through the Tourette Association of America mailing list as well as from 4 TS specialty clinics in the United States. Individuals with a history of intellectual disability, seizure disorder, or a known tic disorder unrelated to TS were excluded.

Additional control subjects were taken from four external large-scale genetic studies consisting of healthy individuals sampled from similar geographic locations and genotyped on the same Illumina OmniExpress platform as the TS cases:

1. Cardiff Controls (CC): UK Blood donors were recruited in Cardiff at the time of blood donation at centers in Wales and England. Although not explicitly screened for psychiatric disorders, these controls are likely to have low rates of severe neuropsychiatric illness, as blood donors in the UK are only eligible to donate if they are not taking any medications. 57% of these controls were male.
2. Consortium for Neuropsychiatric Phenomics (CNP): A collection of neuropsychiatric samples composed of patients with attention deficit hyperactivity disorder (ADHD), bipolar disorder (BD), schizophrenia (SCZ), and psychologically normal controls, collected throughout North America as part of a large NIH Roadmap interdisciplinary research consortia centered at the University of California, Los Angeles. Only the control samples were used in this study.
3. Genomic Psychiatry Cohort (GPC)^28^: A large, longitudinal, population resource composed of clinically ascertained patients affected with BD, SCZ, their unaffected family members, and a large set of control samples with no family history of either disorder. Samples were collected at various sites throughout North America in a National Institute of Mental Health-sponsored study lead by the University of Southern California.
4. Welcome Trust Case Control Consortium 2 (WTCCC2)^29^: Selected control samples from the National Blood Donors Cohort.

### Genotyping and data processing

Cases and controls collected specifically for this study were all genotyped on the Illumina OmniExpress Exome v1.1, while the remaining control samples from the CC, CNP, GPC, and WTCCC2 cohorts were genotyped on the Illumina OmniExpress 12v1.0. The content of these two arrays is identical except for 1) exome-focused content on the former and 2) additional intensity-only markers on the latter. We have observed that exome-specific assays in general exhibit a much higher variance overall in their derived log-R ratio (LRR). Therefore, in order to avoid detection biases due to this differential variance, as well as from unequal probe coverage, only the SNP assays common to all versions of the OmniExpress arrays in this study were used for quality control (QC) and CNV detection, a total of 689,077 markers.

To ensure the generation of the most reliable SNP calls, intensity measures, and B-allele frequencies (BAF), as well as to reduce the effect of differential processing, a custom cluster file was generated for each individual genotyping batch. Since the performance of Illumina’s proprietary normalization and cluster generation process is dependent on the number of samples, we processed all of the raw data, regardless of phenotype, with subsequent removal of clinical samples from the CNP and GPC datasets prior to analysis. An initial round of quality control (QC) was carried out using Illumina Beeline to determine baseline calling rates for each sample using the canonical cluster file (*.egt) provided by the manufacturer for each array version. Any sample with a call rate < 0.98 or a log-R ratio (LRR) standard deviation > 0.30 was deemed a failed assay and removed (Pre-cluster QC, Supplementary Table 1). SNP clustering was then performed in GenomeStudio with only passing samples within each genotyping batch. This process was repeated for all datasets.

### Genome-wide detection of CNV loci

We employed two widely-used HMM-based CNV calling algorithms, PennCNV (version 201105-03) and QuantiSNP (version 2.0), to initially detect structural variants in our dataset. We created GC-adjusted LRR intensity files for all samples using the GC-waviness correction method described by Diskin et al^30^. For PennCNV, a custom population B-allele frequency file was created for each genotyping batch separately and CNV calls were generated using the standard protocol. QuantiSNP calls were generated on the GC-adjusted intensity files. A concordant callset between both CNV callers was then generated by taking the intersecting boundaries of overlapping calls of the same CNV type (deletion or duplication). Additionally, adjacent CNV calls were merged if they were spanned by a CNV called by the other HMM algorithm. As HMMs have been shown to artificially break up large CNVs, we also merged CNV segments in the final concordant callset if they were of the same copy number and the number of intervening markers between them was less than 20% of the total of both segments combined. We repeated this process iteratively until no more joining occurred.

### Sensitivity assessment

Since the samples in this study were consolidated from multiple studies, subtle differences in the ability to detect genetic variation between cases and controls could lead to spurious associations. This issue is even more critical in studies involving CNVs, given the inherent imprecise nature of their detection.

Therefore, prior to association testing, we augmented a previously described method^7^ to investigate whether any difference in sensitivity to detect CNVs existed between cases and controls within the context of our study. Both HMM-based CNV callers we employed for genome-wide detection are univariate methods completely agnostic of intensity information across multiple samples and do not use known population frequency prior probabilities in their calling algorithms. Therefore, common CNVs act as an ideal proxy to evaluate the effectiveness to detect rare events accurately, given that they are detected in the same manner but are present at much higher frequencies.

To facilitate data processing and visualization, we first generated an HDF5 database containing the LogR-Ratio (LRR) intensity and B-allele frequency (BAF) values for all samples. Normalized intensity values across each individual were generated by converting the GC-corrected, median-centered LRR measures into Z-scores. We used the UCSC Genome Browser liftOver tool to translate a list of common HapMap3 CNVs to the hg19 reference. To match the thresholds used for our association tests in this study, we filtered the list of common CNVs to those that were >50 kbp in length. We reduced the number of markers required slightly to a minimum of 9 to ensure that an adequate number of events could be assessed. For each common CNV meeting these criteria, we examined the distribution of median-summarized normalized intensity measures within the CNV region across all study samples and retained only those loci that exhibited discrete clustering into distinct copy-number states. A total of 11 common CNV loci were retained for sensitivity analysis.

We generated locus-specific genotyping calls in the following manner. First, we extracted the LRR intensity Z-scores for all probes in the region across all samples. The Z-scores for all probes spanning the CNV locus were then subjected to a second round of normalization in order to normalize scores across all samples; this was found to aid in the automation of the clustering procedure. A Gaussian mixture model (GMM) was fit to this distribution to cluster samples into discrete CNV groups using the SciKit-learn Python package. The optimum number of clusters was automatically determined by minimization of the Bayesian Information Criterion (BIC) and corrected, when necessary, by manual inspection. Individuals were assigned to a cluster only if the posterior probability of assignment exceeded 0.95.

Copy number state was inferred by examining the original LRR intensity values for samples within each cluster. We inspected for allele frequency differences between controls and cases for all clusters and found no significant difference (Fisher exact test, Supplementary Table 3). We collapsed the clusters at each locus into CNVs of the same type (deletion or duplication). As this locus-specific genotyping method is more sensitive than HMM segmentation methods, we used the proportion of concordant of HMM-based calls as a measure of segmentation detection sensitivity. We found no significant difference in sensitivity to detect common CNVs between phenotypic groups at any of the 11 loci tested, either independently, or in concert (Fisher exact test, Supplementary Tables 4a and 4b). Furthermore, the mean sensitivity for each sample was calculated and collectively assessed for any systematic difference between phenotypic groups. Considering duplications, deletions, or both in concert, we observed no significant difference in the sensitivity of segmentation calls between case and control groups (Welch *t*-test, Supplementary Table 4c), thus validating our preprocessing, QC, and CNV-calling procedures. Results obtained by fitting mixture models separately by either phenotype or batch produced similar results.

### Call filtering, delineation of rare events, and in-silico validation

Calls were removed from the dataset if they spanned less than 10 markers, were less than 50kb in length, or overlapped by more than 0.5 of their total length with regions known to generate artifacts in SNP-based detection of CNVs, including immunoglobulin, telomeric (defined as 100kb from the chromosome ends) and centromeric regions, segmental duplications, and regions that have previously demonstrated associations specific to Epstein-Barr virus immortalized cell lines^29^. We filtered our callset for rare CNVs, defined as those events with MAF approximately < 1% (no more than 65 occurrences across 6,435 samples), based on a reciprocal overlap of 50% with CNVs of the same copy-number state. As the number of rare CNVs in a cohort such as ours is exceedingly large, traditional methods that rely on manual inspection of all putative CNV calls constitutes an approach that is both inconsistent and impractical. Therefore, for each putative rare CNV, we generated two different metrics based on intensity (LRR-Z) and BAF banding (BAF_del_ and BAF_dup_), and calculated population frequency estimates (OUTLIER-Z) by inspecting the distribution of intensities across the entire sample (Supplementary Figure 4a).

For qualifying CNVs based on intensity, we adopted a scoring methodology similar to the MeZOD method described previously^32,33^, with a noted exception. We observed that standardized intensity measures from Illumina data typically range from < −20 for homozygous deletions, [−6,−2.5] for heterozygous deletions, and > 1.5 for duplications. Because of the disproportionately large effect on intensity measures caused by deletion events, performing a second round of normalization across all samples within each putative CNV often skews the overall distribution when such events are present. Therefore, we only performed a single round of normalization of LRR intensity measures within each sample. Each CNV is scored by calculating the median of LRR intensity Z-scores (LRR-Z) for all probes within the region. To determine reasonable thresholds for intensity metrics specific to our Omniexpress assay, we applied our CNV calling pipeline on 266 HapMap samples genotyped on the same OmniExpress platform, provided by Illumina. We compared these HapMap calls to those generated using high-coverage array comparative genome hybridization (aCGH)^34^. 11 samples overlapped between these two datasets. For these individuals, we extracted the LRR-Z for all aCGH-validated CNV calls and inspected the distribution of all calls for both deletions and duplications in order to establish reasonable, conservative thresholds based on validated CNVs; in this case we required an LRR-Z of <−2.3 for deletions and >+1.3 for duplications (Supplementary Figure 4b).

The BAF banding pattern is particularly informative for the detection of CNV events using SNP arrays. This is particularly true for duplications, as intensity gains are typically modest for these types of events. We calculated the proportion of probes within the CNV region that showed evidence of a deletion (BAF of <0.15 or >0.85) or duplication event (BAF of [0.25-04] or [0.60.8]), and denoted these measures “BAF_del_” and “BAF_dup_”. Based on prior observations^35^ and from examination of our own data, we conservatively required deletions to have BAF_del_ > 0.9, and duplications to have a BAF_dup_ > 0.15.

Furthermore, the distribution of summarized intensity information across all individuals for every putative rare CNV was screened for calling sensitivity by inspecting the distribution of intensities at the locus across all samples. For each rare CNV, we flagged those where the proportion of samples whose LRR-Z metric fell outside of [−2.3, 1.3] (denoted as OUTLIER-Z) and further inspected these regions manually. Putatively rare CNV loci that showed substantial evidence for extensive polymorphism were subsequently scored using the GMM genotyping method described above.

We opted not to impose any hard cutoff for CNV calls with regard to any of these measures to avoid any bias imposed by differential missingness derived from subtle systematic differences between genotyping batches. Rather, these thresholds were applied to flag those CNV calls with marginal scores for manual inspection and filter out only obviously misclassified events. Through this *in silico* validation process, we discovered multiple instances of large copy-number aberrations likely due to individual mosaicism (Supplementary Figure 4c), and-two common polymorphic duplication regions misclassified as a rare CNV due to reduced sensitivity of the HMM segmentation (Supplementary Figure 4d). Out of 8,452 initial consensus HMM calls, a total of 87 CNV events were removed. Six of these were due to mosaic events, and the remainder was excluded due to the misclassification of common CNVs as a rare events.

### Genome-wide burden analysis

For comparison of genome-wide burden between TS cases and controls, we limited our consideration to rare CNVs spanning a minimum of 10 SNPs and > 50kb in length. We assessed genome-wide CNV burden in three different ways: (1) number of rare CNVs, (2) total CNV length (per 100 KB), and (3) the total number of genes intersected by CNVs. Furthermore, we stratified each test by both size (all CNVs, >500kb, and >1Mb) as well as frequency (rare CNVs and singletons). Frequency counts were determined using PLINK `--cnv-freq-method2 0.5.` Here, singletons are defined as sharing no more than a 50% overlap with any other CNV. Gene overlaps were counted if the CNV overlapped any gene boundary as delineated by Refseq. To examine the effect of different covariates on our different metrics of global CNV burden, we first fit a linear regression model for each type of burden test:

> *Burden*_*metric* ~ *genotyping*_*batch* + *subject*_*sex* + *LRR*_*SD* + *ancestry*_*PCs*

None of the included covariates, which included the top 10 ancestry PCs, were significant predictors of global CNV burden as measured by either total number of CNVs or cumulative CNV length (Supplementary Table 6a). Separately, we examined the effect of these covariates exclusively with regard to the burden due to small CNV events, as these are most likely affected by minor fluctuations in assay quality and subtle differences in sample ascertainment. We found that LRR_SD was significantly associated with small CNV burden (defined here as CNVs < 100kb in length) for both total CNV number and CNV size (Supplementary Table 6b and 6c), and was therefore included in the burden analysis as a covariate.

To assess for a global burden difference between TS and controls, we fit a logistic regression model in R with affectation status as the dependent variable and the burden metric and LRR_SD as independent variables. ORs indicate an increase in risk for TS as assessed per CNV, per 100kb of total CNV length, or per gene affected by CNVs. P-values were calculated using the likelihood ratio test.

### Validation of association results

We repeated the segmental association test after carefully pair-matching each case with a control such that the global difference between each pair was minimized using SpectralGEM^32^ (Supplementary Figure 8). For the matched segmental association analysis, because of the drastic reduction in sample size, a corrected p-value < 0.05 was used as a cutoff to indicate genome-wide significance.

**Supplementary Figure 1.**
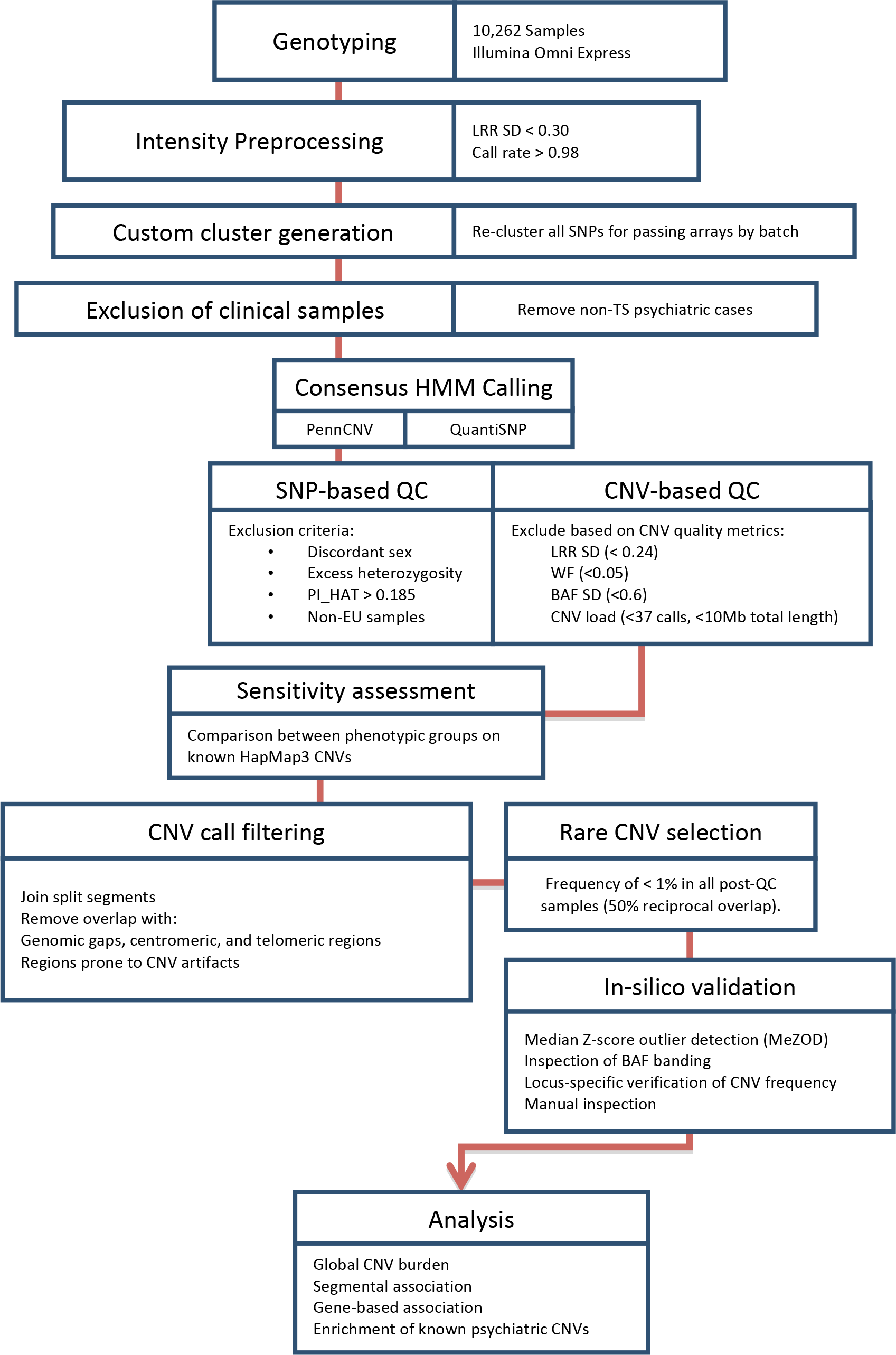
Overview of the CNV calling pipeline and analysis. A schematic of the data processing, CNV calling, and analysis is presented here, and described in detail in the supplementary text.

**Supplementary Table 1.** (a) Summary of included studies and genotyping information. Sample phenotypes, genotyping platform, and genotyping center for different datasets collected for this study are shown, separated by study. (b) Summary of quality control procedures by study. The number of samples remaining within each batch after each successive quality control step (as described in the Supplementary Text) is shown. Study abbreviations: Cardiff Controls (CC), Consortium for Neuropsychiatric Phenomics (CNP), Genomic Psychiatry Cohort (GPC), Wellcome Trust Case-Control Consortium (WTCCC_2_) and TS cases and controls collected for this study (TS1-3).

| Supplementary Table 1a: Studies and genotyping |  |  |  |
| --- | --- | --- | --- |
| GENOTYPING BATCH | ARRAY | CENTER | PHENOTYPES |
| **CC** | OmniExpress v 1.0 | Cardiff | Control |
| **CNP** | OmniExpress v 1.0 | Broad | Control/Clinical |
| **GPC** | OmniExpress v 1.0 | Broad | Control/Clinical |
| **WTCCC2** | OmniExpress v 1.0 | Cardiff | Control |
| **TS1** | OmniExpress Exome v 1.1 | UCLA | Control/TS |
| **TS2** | OmniExpress Exome v 1.1 | UCLA | Control/TS |
| **TS3** | OmniExpress Exome v 1.1 | UCLA | Control/TS |

| Supplementary Table 1b: QC summary |  |  |  |  |  |  |  |  |
| --- | --- | --- | --- | --- | --- | --- | --- | --- |
| QC STEP | CC | CNP | GPC | WTCCC2 | TS1 | TS2 | TS3 | TOTALS |
| **Initial Samples** | 1,146 | 1,511 | 3,197 | 960 | 1,152 | 2,160 | 136 | 10,262 |
| **Pre-cluster QC** | 1,141 | 1,510 | 3,126 | 870 | 1,148 | 2,152 | 135 | 10,082 |
| **Duplicate/Control Samples** | 1,141 | 1,491 | 3,081 | 870 | 1,134 | 2,143 | 134 | 9,994 |
| **Clinical Phenotype** | 1,141 | 1,312 | 1,388 | 870 | 1,134 | 2,143 | 134 | 8,122 |
| **Sex Concordance** | 1,141 | 1,312 | 1,387 | 870 | 1,132 | 2,140 | 134 | 8,116 |
| **Heterozygosity** | 1,122 | 1,247 | 1,301 | 859 | 1,107 | 2,089 | 133 | 7,858 |
| **Cryptic Relatedness** | 1,084 | 1,222 | 1,264 | 852 | 1,100 | 2,067 | 132 | 7,621 |
| **CNV/Intensity QC** | 991 | 1,106 | 1,164 | 832 | 1,014 | 1,843 | 119 | 6,928 |
| **EU Ancestry** | 959 | 572 | 1,143 | 813 | 959 | 1,803 | 116 | 6,535 |

**Supplementary Figure 2.**
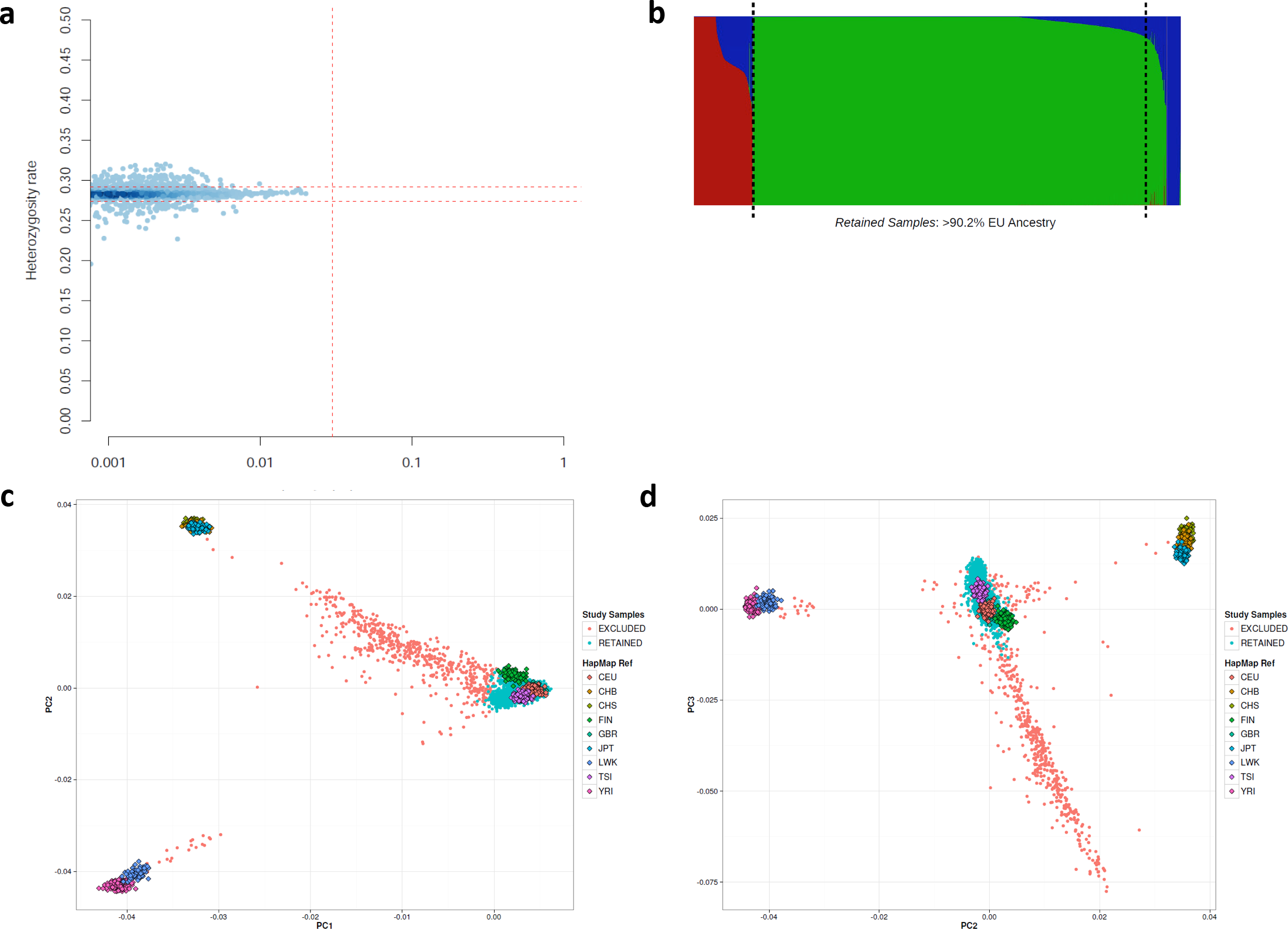
SNP-based quality control and ancestry determination. (a) Exclusion of sample outliers based on heterozygosity, mean +/− 1.5 SD (red dotted lines). (b) Exclusion of non-European samples based on ethnicity estimation using fastStructure with HapMap continental groups and K=3 clustering. Samples with > 9.85% non-EU ancestry were excluded. This threshold was calibrated against the maximum of reference European groups CEU, GBR, and TSI. The results of principal component (PC) analysis for the cohort and reference groups are plotted along (c) PCs 1 and 2 and (d) PCs 2 and 3. Retained samples and excluded samples are shown in cyan and pink, respectively.

**Supplementary Figure 3.**
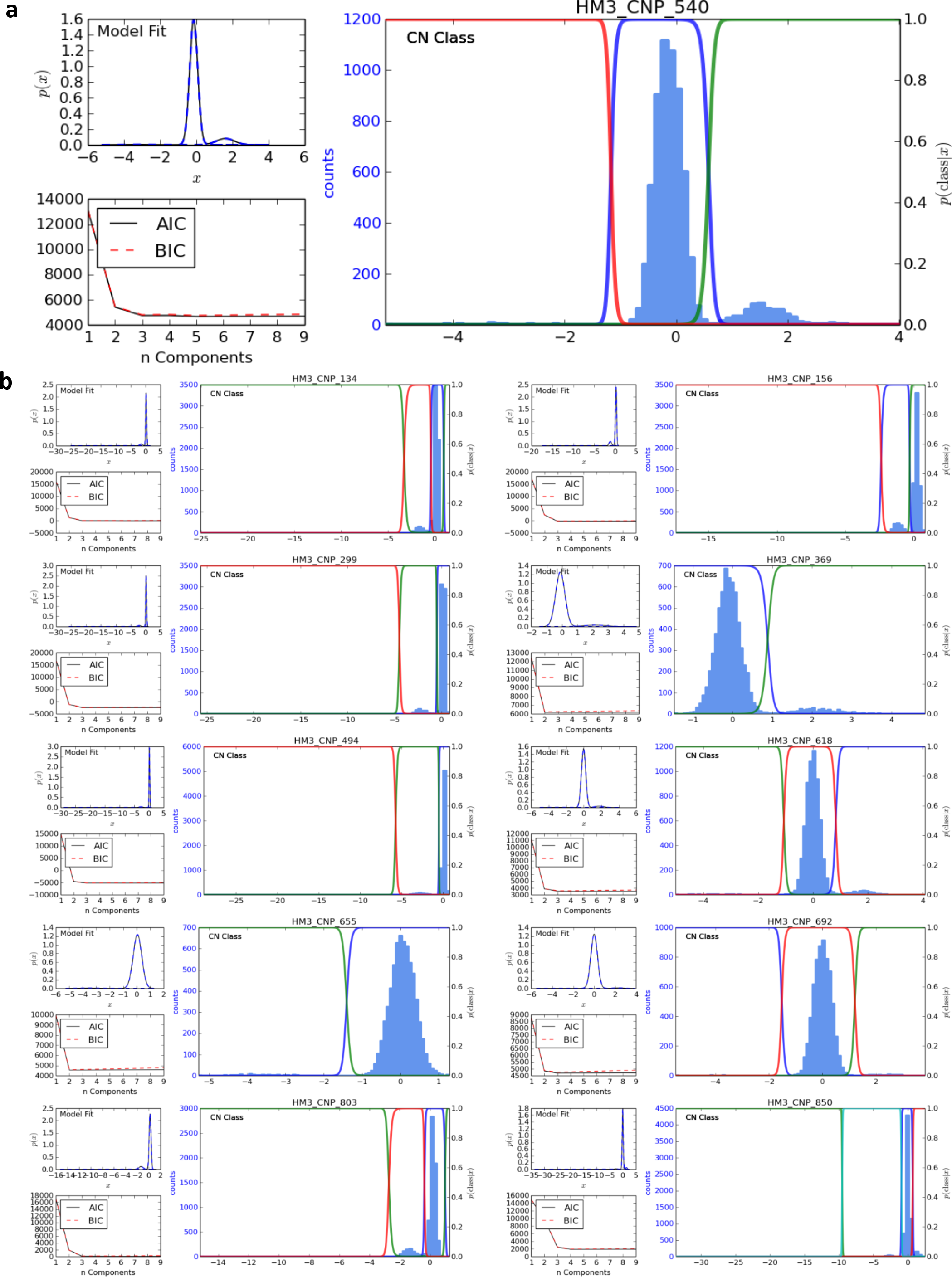
Gaussian mixture model clusters of common HM3 CNVs. (a) A representative GMM cluster plot for locus HM3_CNP_540. Subplots for each CNV depict, counter-clockwise: the best-fit model, Akaike and Bayesian Information Criterion metrics calculated for GMM fitting 1-9 components, and the posterior probability for CNV cluster assignment (colored lines) overlaying the distribution of median summarized intensity values for all samples across region calculated using the best-fit model. (b) GMM plots for the 10 additional HapMap3 CNV loci that were used to critically evaluate sensitivity between cases and controls are displayed on the following page (Supplementary Methods and Supplementary Table 3)

**Supplementary Table 3.** Gaussian Mixture Model (GMM) clustered genotype calls at common Hapmap 3 CNVs. For sensitivity analysis, all 6,535 samples used in this study were genotyped across 11 common Hapmap3 CNVs using a locus-specific GMM-based clustering method (see Supplementary Methods). The numbers for CNV_ID: Hapmap3 accession number. CLUSTER_ID: Arbitrary identifier assigned by the clustering algorithm. CLUSTER_LRR: The mean value of all median-summarized intensity values for all samples assigned to the cluster. CLUSTER_COPY: Copy-number state inferred by examination of raw LRR-intensity values for samples within the cluster. Call frequencies (FREQ) for 4,100 controls (CTRL) and 2,435 TS cases (CASE) reflect the proportion of GMM-based genotype calls with > 0.95 posterior probability of cluster assignment. There was no significant difference in CNV genotype frequency between phenotypic groups at any of the 21 non-reference genotype calls across all 11 loci (Fisher’s exact test).

| Supplementary Table 3: GMM-based genotype calls at common CNV loci |  |  |  |  |  |  |  |  |
| --- | --- | --- | --- | --- | --- | --- | --- | --- |
| CNV_ID | CLUSTER | CLUSTER_LRR | GMM_COPY | CTRL_CALLS | CASE_CALLS | CTRL_FREQ | CASE_FREQ | p-value |
| HM3_CNP_134 | 1 | -13.17478812 | 0 | 7 | 4 | 0.002 | 0.002 | 1.0 |
| HM3_CNP_134 | 2 | -1.544234008 | 1 | 296 | 191 | 0.072 | 0.078 | 0.4 |
| HM3_CNP_156 | 1 | -1.141264855 | 1 | 517 | 315 | 0.126 | 0.129 | 0.7 |
| HM3_CNP_156 | 2 | -10.96495207 | 0 | 18 | 13 | 0.004 | 0.005 | 0.6 |
| HM3_CNP_299 | 1 | -2.148959932 | 1 | 275 | 142 | 0.067 | 0.058 | 0.2 |
| HM3_CNP_299 | 2 | -20.27406193 | 0 | 4 | 3 | 0.001 | 0.001 | 0.7 |
| HM3_CNP_369 | 1 | 2.201328191 | 3 | 234 | 137 | 0.057 | 0.056 | 0.9 |
| HM3_CNP_494 | 1 | -2.7402128 | 1 | 196 | 100 | 0.048 | 0.041 | 0.2 |
| HM3_CNP_494 | 2 | -23.74078464 | 0 | 1 | 1 | 0.000 | 0.000 | 1.0 |
| HM3_CNP_540 | 1 | 1.648736036 | 3 | 392 | 263 | 0.096 | 0.108 | 0.1 |
| HM3_CNP_540 | 2 | -3.301784945 | 1 | 32 | 17 | 0.008 | 0.007 | 0.8 |
| HM3_CNP_618 | 1 | -3.470922513 | 1 | 44 | 24 | 0.011 | 0.010 | 0.8 |
| HM3_CNP_618 | 2 | 1.817743156 | 3 | 167 | 91 | 0.041 | 0.037 | 0.5 |
| HM3_CNP_655 | 1 | -3.645609914 | 1 | 45 | 34 | 0.011 | 0.014 | 0.3 |
| HM3_CNP_692 | 0 | -4.175170396 | 1 | 10 | 11 | 0.002 | 0.005 | 0.2 |
| HM3_CNP_692 | 2 | 2.34262532 | 3 | 47 | 32 | 0.011 | 0.013 | 0.6 |
| HM3_CNP_803 | 1 | -10.64502833 | 0 | 15 | 14 | 0.004 | 0.006 | 0.2 |
| HM3_CNP_803 | 2 | -1.347106673 | 1 | 417 | 258 | 0.102 | 0.106 | 0.6 |
| HM3_CNP_850 | 1 | -31.82000658 | 0 | 1 | 1 | 0.000 | 0.000 | 1.0 |
| HM3_CNP_850 | 2 | -2.649145422 | 1 | 74 | 49 | 0.018 | 0.020 | 0.6 |
| HM3_CNP_850 | 3 | 1.410233747 | 3 | 165 | 101 | 0.040 | 0.041 | 0.8 |

**Supplementary Table 4.**
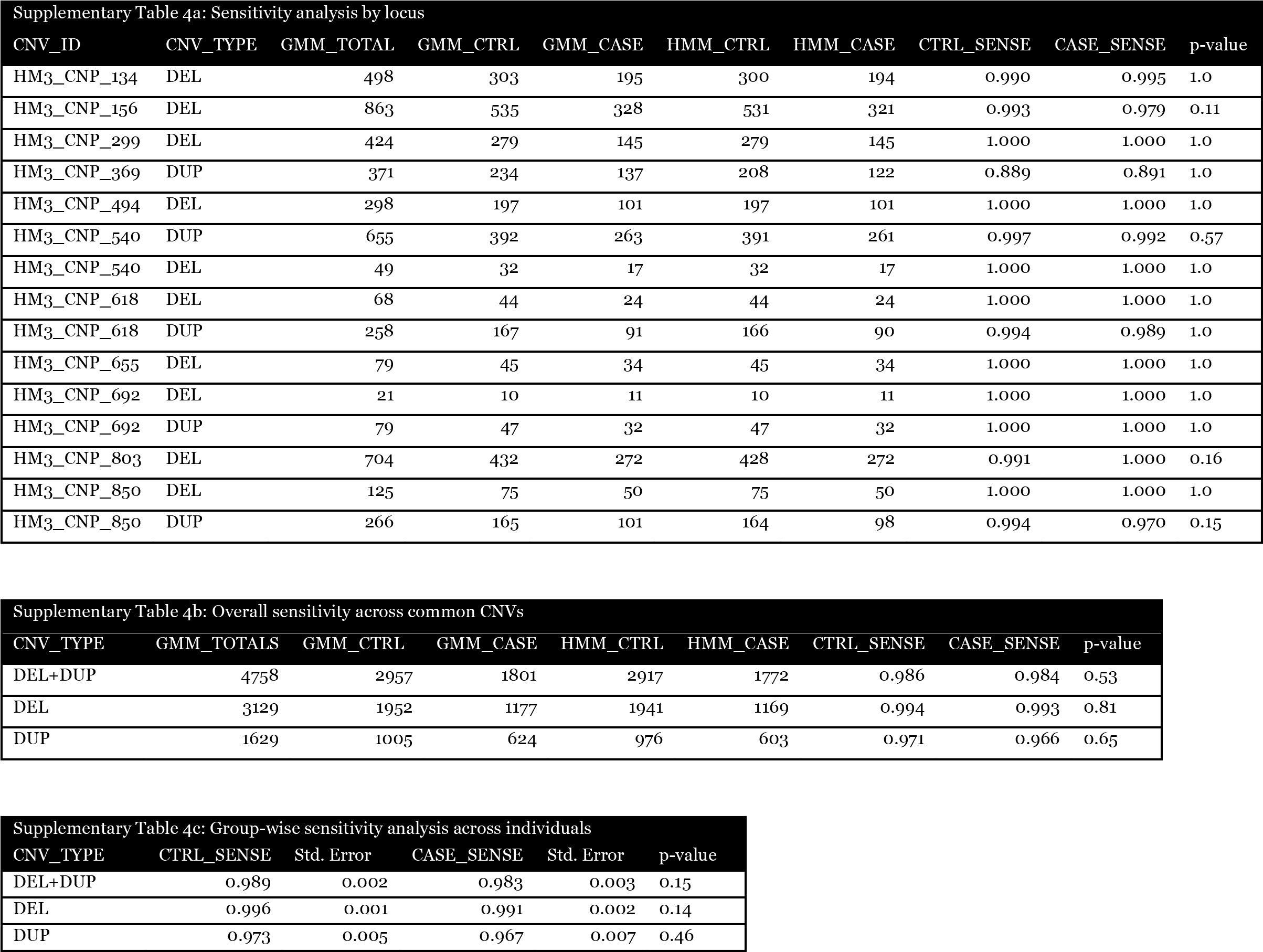
Sensitivity analysis of consensus HMM-segmentation calls. (a) Sensitivity by locus. The sensitivity HMM calling was defined as the number of concordant HMM calls divided by the total number of non-reference genotypes called in the same individual by GMM clustering. Non-reference GMM calls were collapsed into calls of the same class (CNV_TYPE, DEL or DUP). (b) Overall sensitivity across all loci. P-values for both individual and aggregated locus-specific locus-based tests were calculated using Fisher’s exact test. (c) Group-wise comparison of sensitivity between cases and controls based on the average sensitivity calculated within each individual. No significant difference in the average individual sensitivity was observed between phenotypic groups whether considering deletions, duplications, or both in concert (Welch’s *t*-test).

| Supplementary Table 4a: Sensitivity analysis by locus |  |  |  |  |  |  |  |  |  |
| --- | --- | --- | --- | --- | --- | --- | --- | --- | --- |
| CNV_ID | CNV_TYPE | GMM_TOTAL | GMM_CTRL | GMM_CASE | HMM_CTRL | HMM_CASE | CTRL_SENSE | CASE_SENSE | p-value |
| HM3_CNP_134 | DEL | 498 | 303 | 195 | 300 | 194 | 0.990 | 0.995 | 1.0 |
| HM3_CNP_156 | DEL | 863 | 535 | 328 | 531 | 321 | 0.993 | 0.979 | 0.11 |
| HM3_CNP_299 | DEL | 424 | 279 | 145 | 279 | 145 | 1.000 | 1.000 | 1.0 |
| HM3_CNP_369 | DUP | 371 | 234 | 137 | 208 | 122 | 0.889 | 0.891 | 1.0 |
| HM3_CNP_494 | DEL | 298 | 197 | 101 | 197 | 101 | 1.000 | 1.000 | 1.0 |
| HM3_CNP_540 | DUP | 655 | 392 | 263 | 391 | 261 | 0.997 | 0.992 | 0.57 |
| HM3_CNP_540 | DEL | 49 | 32 | 17 | 32 | 17 | 1.000 | 1.000 | 1.0 |
| HM3_CNP_618 | DEL | 68 | 44 | 24 | 44 | 24 | 1.000 | 1.000 | 1.0 |
| HM3_CNP_618 | DUP | 258 | 167 | 91 | 166 | 90 | 0.994 | 0.989 | 1.0 |
| HM3_CNP_655 | DEL | 79 | 45 | 34 | 45 | 34 | 1.000 | 1.000 | 1.0 |
| HM3_CNP_692 | DEL | 21 | 10 | 11 | 10 | 11 | 1.000 | 1.000 | 1.0 |
| HM3_CNP_692 | DUP | 79 | 47 | 32 | 47 | 32 | 1.000 | 1.000 | 1.0 |
| HM3_CNP_803 | DEL | 704 | 432 | 272 | 428 | 272 | 0.991 | 1.000 | 0.16 |
| HM3_CNP_850 | DEL | 125 | 75 | 50 | 75 | 50 | 1.000 | 1.000 | 1.0 |
| HM3_CNP_850 | DUP | 266 | 165 | 101 | 164 | 98 | 0.994 | 0.970 | 0.15 |

| Supplementary Table 4b: Overall sensitivity across common CNVs |  |  |  |  |  |  |  |  |
| --- | --- | --- | --- | --- | --- | --- | --- | --- |
| CNV_TYPE | GMM_TOTALS | GMM_CTRL | GMM_CASE | HMM_CTRL | HMM_CASE | CTRL_SENSE | CASE_SENSE | p-value |
| DEL+DUP | 4758 | 2957 | 1801 | 2917 | 1772 | 0.986 | 0.984 | 0.53 |
| DEL | 3129 | 1952 | 1177 | 1941 | 1169 | 0.994 | 0.993 | 0.81 |
| DUP | 1629 | 1005 | 624 | 976 | 603 | 0.971 | 0.966 | 0.65 |

| Supplementary Table 4c: Group-wise sensitivity analysis across individuals |  |  |  |  |  |
| --- | --- | --- | --- | --- | --- |
| CNV_TYPE | CTRL_SENSE | Std. Error | CASE_SENSE | Std. Error | p-value |
| DEL+DUP | 0.989 | 0.002 | 0.983 | 0.003 | 0.15 |
| DEL | 0.996 | 0.001 | 0.991 | 0.002 | 0.14 |
| DUP | 0.973 | 0.005 | 0.967 | 0.007 | 0.46 |

**Supplementary Figure 4.**
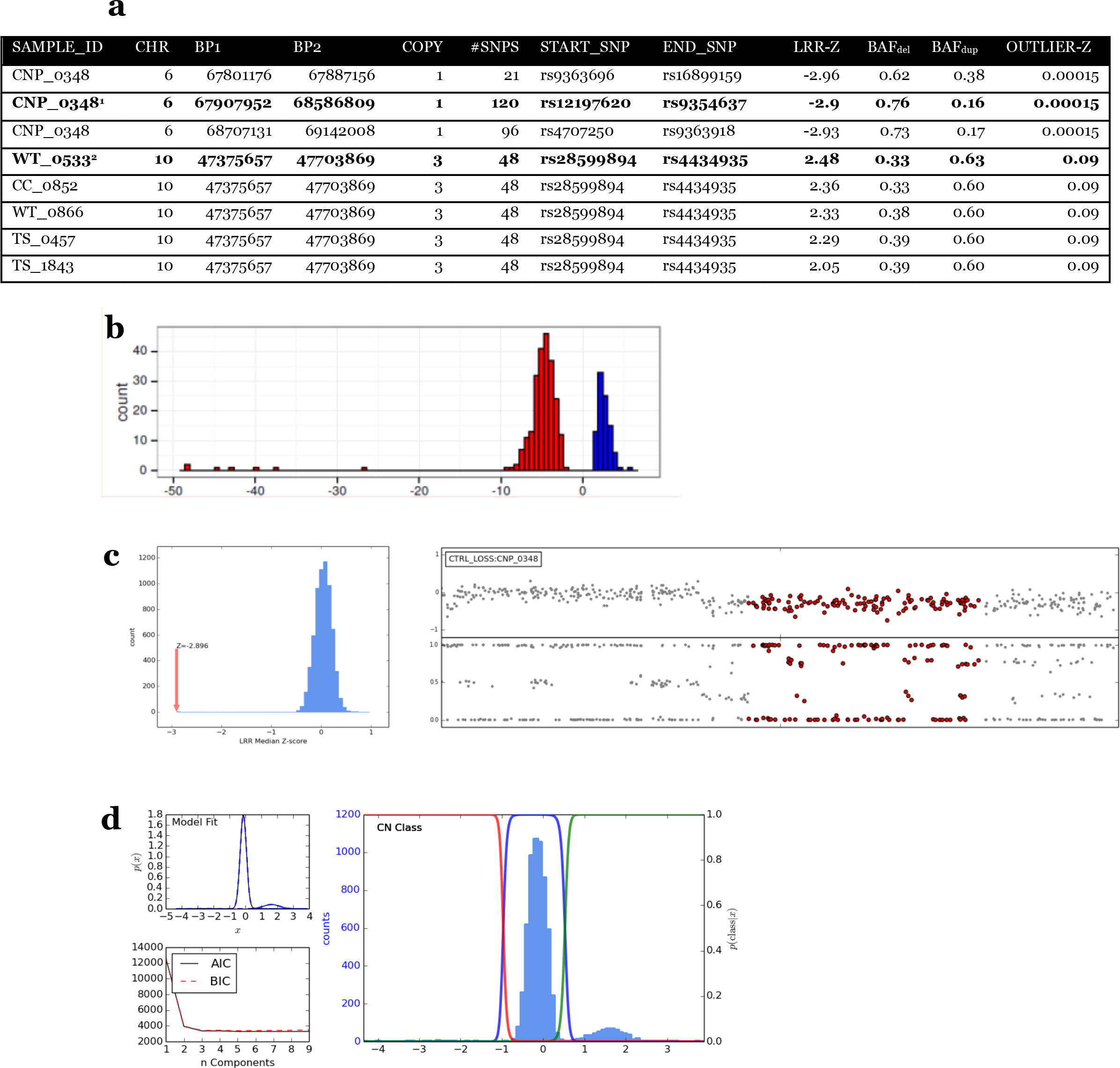
In-silico validation of CNV calls. (a) Representative CNVs scored with various CNV validation metrics. Abbreviations (see Supplementary Methods for details): median summarized intensity measures across a putative CNV locus, standardized by sample (LRR-Z), proportion of probes with a BAF banding pattern indicative of a duplication event (BAF-D), proportion of samples with LRR-Z scores indicative of a polymorphic event (OUTLIER-Z). (b) Distribution of median summarized standardized intensity values (LRR-Z) for validated CNVs derived from HapMap samples generated on identical arrays as used in this study (Illumina OmniExpress). Based on this distribution, CNVs without an LRR-Z score < 2.3 (deletions) and > 1.3 (duplications) were flagged for manual inspection. (c) Example of a large singleton mosaic event flagged for exclusion in sample CNP_0348, indicated as (1) in Figure 4a. This CNV on chromosome 6 was detected as three separate CNVs after taking the consensus of two different HMM calling algorithms. The largest CNV call exhibits an LRR-Z score of −2.86 (left, red arrow), indicative of a deletion, but shows a clear BAF-banding pattern of a duplication event (right), with a BA_dup_ score of 0.16. This is indicative of a mosaic event, where only a proportion of cells from sample CNP_0348 harbor the deletion event. (d) Example of a polymorphic CNV on chr10:47,375,657-47,703,869 misclassified as a rare event due to reduced sensitivity, indicated as (2) in Figure 4a, with an OUTLIER-Z score of 0.09. Genotyping using GMM-based clustering indicated that this misclassified rare event (MAF < 0.01) has a MAF of 0.12.

**Supplementary Table 5.** Analysis of global CNV burden. Comparison of genome-wide CNV burden among TS cases compared to controls, as defined by (a) total number of CNVs (b) 100-kbp of CNV length, and (c) the total number of genes affected by CNVs, stratified by CNV frequency, size and type. Odds ratios indicate an increased risk for TS per unit of CNV burden, and were calculated using logistic regression with covariates significantly associated with any of the CNV burden metrics used, both for total CNV burden as well as burden due to small CNV events (see Supplementary Methods). P-values were calculated using the likelihood ratio test. Singletons are defined as CNVs occurring only once among either a single case or control. Event counts are based on a 50% reciprocal overlap. Gene counts are defined as the number of Refseq genes overlapping CNVs by any amount.

| Supplementary Table 5a: Burden per CNV |  |  |  |  |  |  |  |
| --- | --- | --- | --- | --- | --- | --- | --- |
|  |  | RARE CNVS |  |  | SINGLETONS |  |  |
| SIZE CATEGORY | CNV TYPE | OR | 95% CI | p-value | OR | 95% CI | p-value |
| ALL | DEL+DUP | 1.01 | [0.97-1.06] | 0.60 | 1.09 | [1.01-1.18] | 0.03 |
|  | DEL | 1.03 | [0.96-1.10] | 0.41 | 1.12 | [0.99-1.26] | 0.06 |
|  | DUP | 1.00 | [0.94-1.06] | 0.97 | 1.07 | [0.96-1.9] | 0.18 |
| >500KB | DEL+DUP | 1.19 | [1.01-1.41] | 0.04 | 1.30 | [1.01-1.67] | 0.04 |
|  | DEL | 1.21 | [0.86-1.71] | 0.26 | 1.72 | [1.06-2.83] | 0.03 |
|  | DUP | 1.18 | [0.98-1.43] | 0.09 | 1.17 | [0.86-1.57] | 0.32 |
| >1MB | DEL+DUP | 1.50 | [1.10-2.03] | 0.009 | 1.91 | [1.24-2.97] | 0.004 |
|  | DEL | 1.85 | [1.06-3.25] | 0.03 | 2.82 | [1.36-6.18] | 0.007 |
|  | DUP | 1.37 | [0.95-1.96] | 0.09 | 1.53 | [0.89-2.64] | 0.12 |

| Supplementary Table 5b: Burden per 100KB of CNV length |  |  |  |  |  |  |  |
| --- | --- | --- | --- | --- | --- | --- | --- |
|  |  | RARE CNVS |  |  | SINGLETONS |  |  |
| SIZE CATEGORY | CNV TYPE | OR | 95% CI | p-value | OR | 95% CI | p-value |
| ALL | DEL+DUP | 1.011 | [1-1.021] | 0.05 | 1.022 | [1.006-1.035] | 0.006 |
|  | DEL | 1.017 | [0.997-1.037] | 0.09 | 1.027 | [1.003-1.055] | 0.04 |
|  | DUP | 1.008 | [0.995-1.021] | 0.22 | 1.016 | [1-1.034] | 0.06 |
| >500KB | DEL+DUP | 1.015 | [1.003-1.027] | 0.01 | 1.021 | [1.006-1.037] | 0.006 |
|  | DEL | 1.016 | [0.994-1.039] | 0.16 | 1.029 | [1-1.059] | 0.06 |
|  | DUP | 1.015 | [1.001-1.029] | 0.04 | 1.017 | [1-1.039] | 0.06 |
| >1MB | DEL+DUP | 1.018 | [1.005-1.032] | 0.007 | 1.024 | [1.007-1.043] | 0.008 |
|  | DEL | 1.025 | [1.001-1.052] | 0.05 | 1.029 | [1.005-1.064] | 0.06 |
|  | DUP | 1.015 | [1-1.032] | 0.05 | 1.021 | [1.002-1.044] | 0.05 |

| Supplementary Table 5c: Burden per gene |  |  |  |  |  |  |  |
| --- | --- | --- | --- | --- | --- | --- | --- |
|  |  | RARE CNVS |  |  | SINGLETONS |  |  |
| SIZE CATEGORY | CNV TYPE | OR | 95% CI | p-value | OR | 95% CI | p-value |
| ALL | DEL+DUP | 1.008 | [1-1.016] | 0.06 | 1.016 | [1.004-1.030] | 0.01 |
|  | DEL | 1.012 | [0.991-1.033] | 0.27 | 1.023 | [0.989-1.059] | 0.18 |
|  | DUP | 1.007 | [0.999-1.016] | 0.10 | 1.015 | [1.002-1.030] | 0.03 |
| >500KB | DEL+DUP | 1.012 | [1.003-1.022] | 0.01 | 1.019 | [1.005-1.036] | 0.03 |
|  | DEL | 1.008 | [0.982-1.034] | 0.53 | 1.024 | [0.984-1.066] | 0.39 |
|  | DUP | 1.012 | [1.002-1.023] | 0.02 | 1.018 | [1.004-1.037] | 0.04 |
| >1MB | DEL+DUP | 1.014 | [1.004-1.026] | 0.01 | 1.022 | [1.008-1.044] | 0.02 |
|  | DEL | 1.021 | [0.989-1.054] | 0.20 | 1.036 | [0.991-1.081] | 0.27 |
|  | DUP | 1.013 | [1.002-1.025] | 0.02 | 1.022 | [1.005-1.044] | 0.04 |

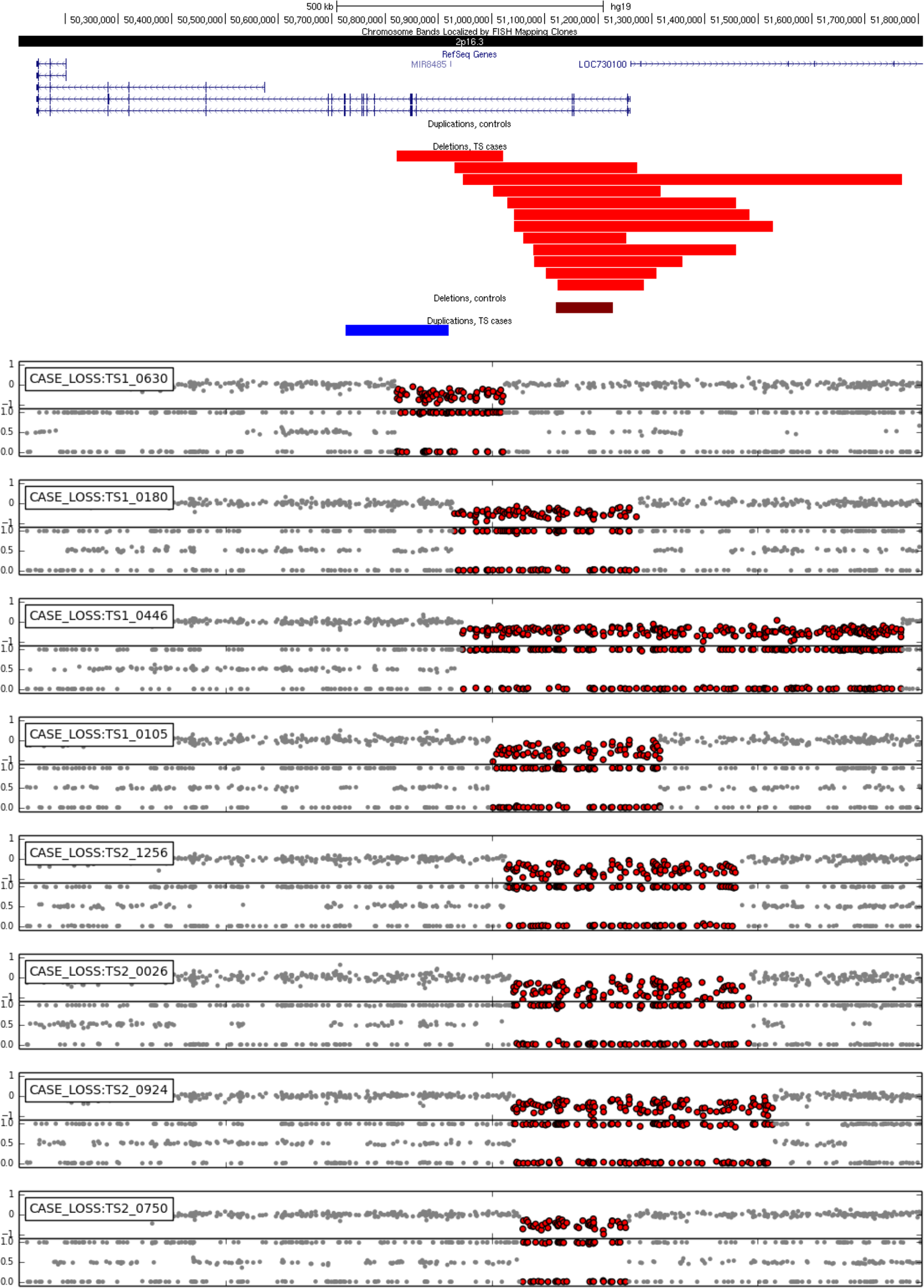

**Supplementary Figure 6.**
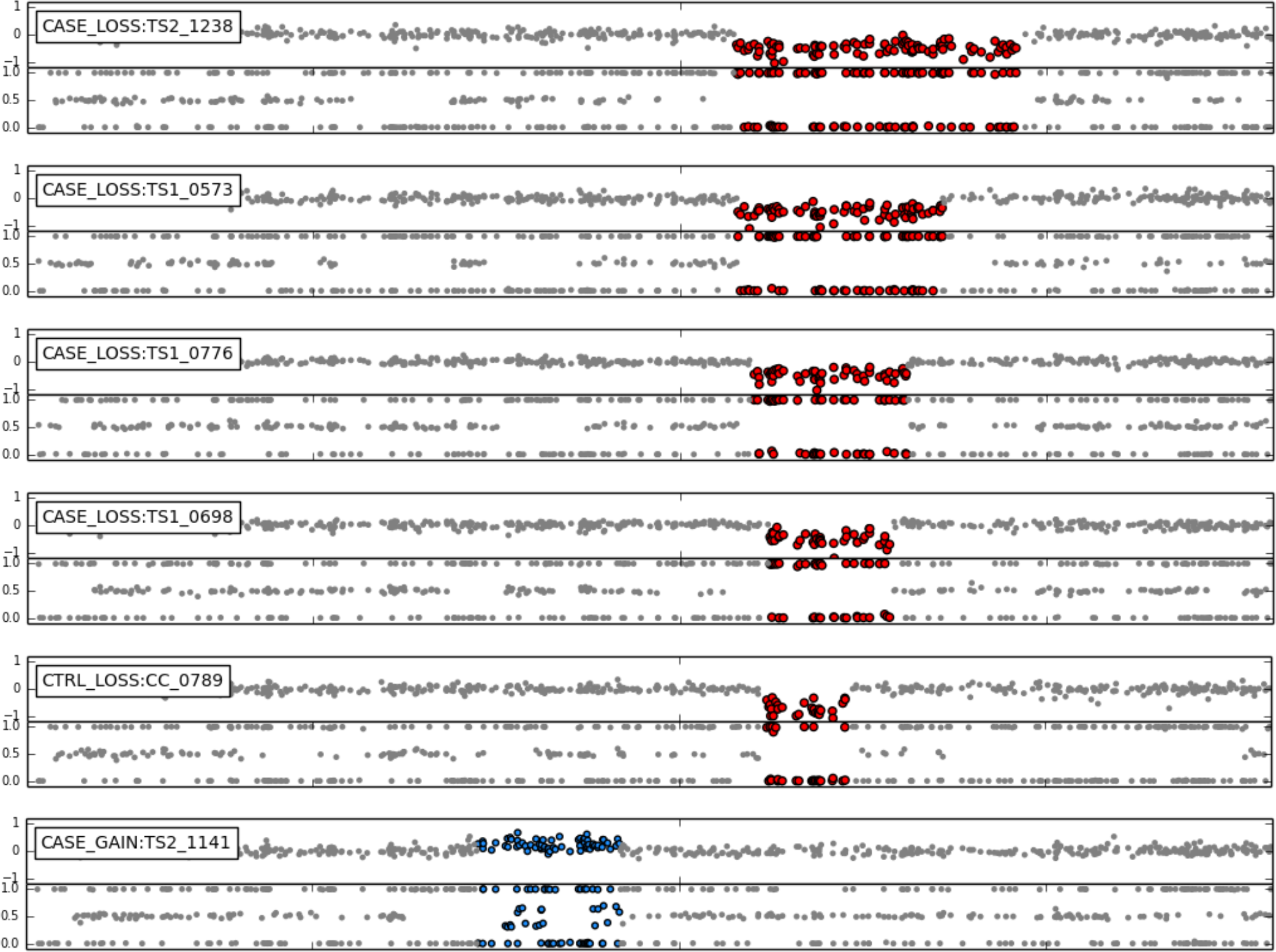
Exonic CNVs affecting NRXN1. UCSC genome browser track depicting all exonic *NRXN_1_* CNVs > 50kb identified in this study: 12 heterozygous case deletions (red), one control deletion (dark red) and a single case duplication (blue). Probe-level plots of LRR intensity and BAF for all exonic *NRXN_1_* CNV carriers shown in the same order as the UCSC genome browser track. Colored probes indicate the location of called deletions (red) and duplications (blue).

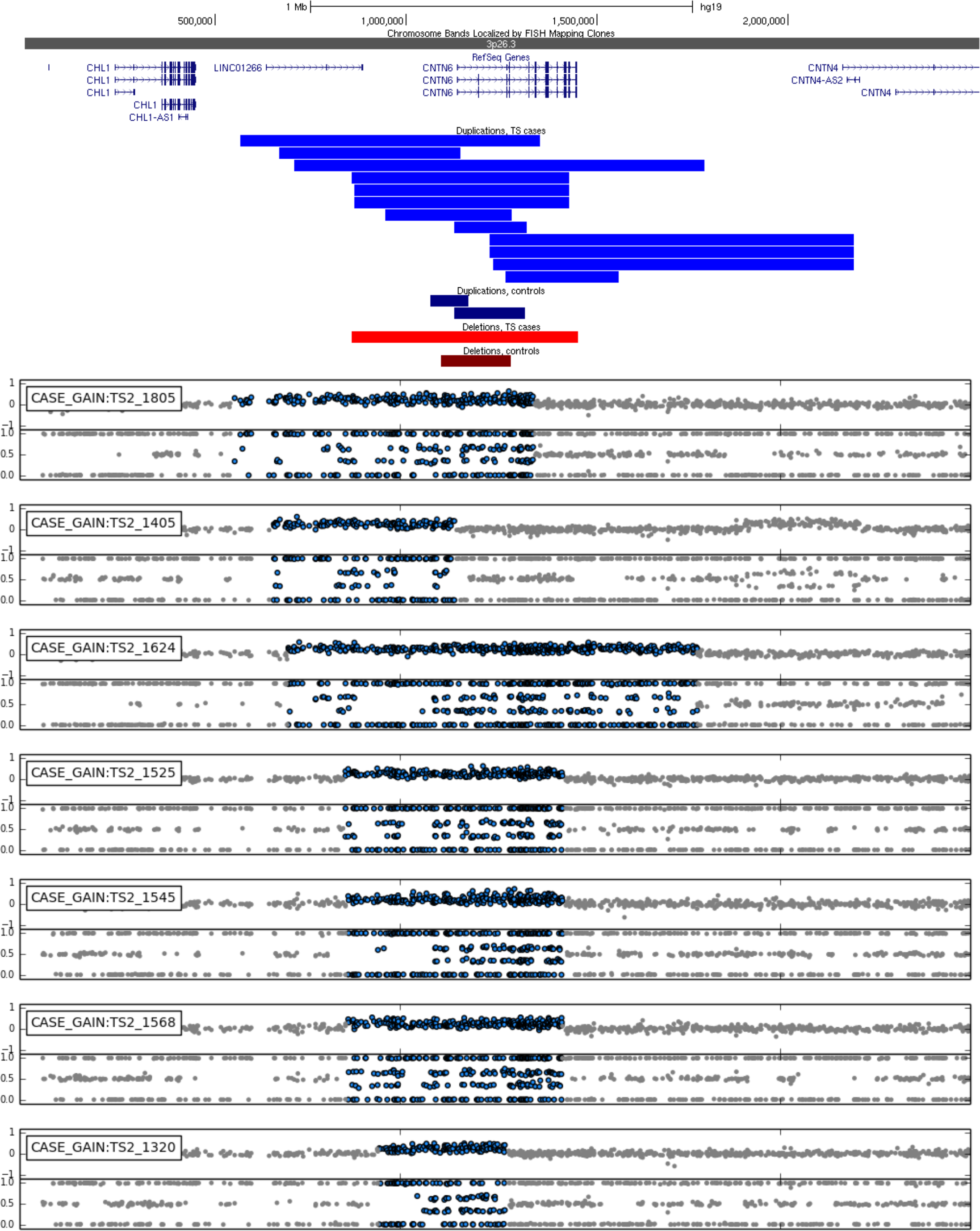

**Supplementary Figure 7.**
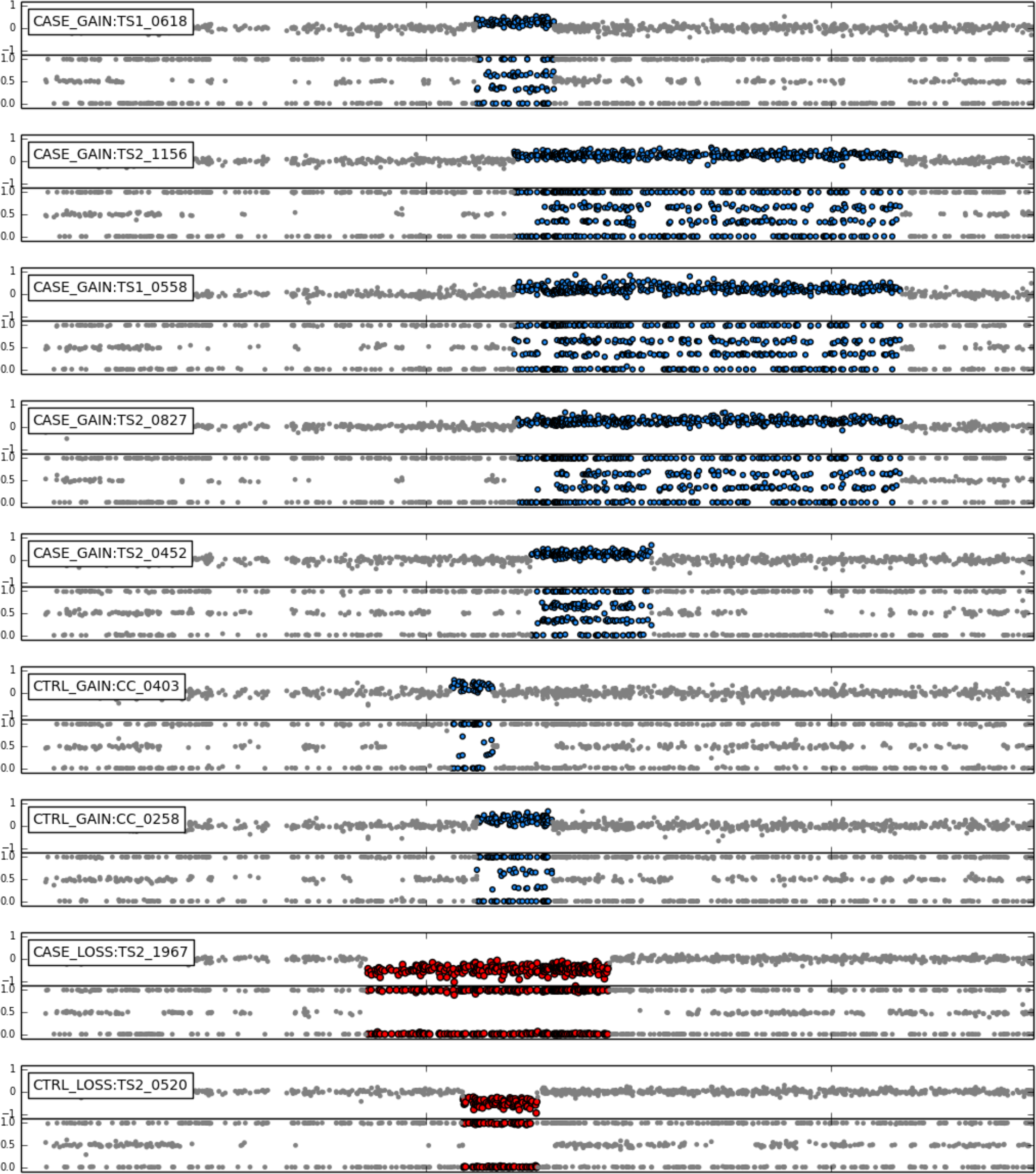
CNVs overlapping *CNTN6*. UCSC genome browser track displaying heterozygous genic duplications in TS cases (blue) and controls (dark blue) followed by deletions (red). Probe-level LRR and BAF plots for all _1_6 CNVs detected spanning *CNTN6* are shown below.

**Supplementary Table 6.** All nominally significant results from gene-based association test. Gene-based association was conditioned on CNVs spanning exons of protein-coding genes, as determined by strict overlap with Refseq annotation. Significance was established using 1,000,000 permutations (P_locus_), using the max(T) method to establish genome-wide significance, corrected for multiple testing (P_corr_). Duplications and deletions were independently filtered for a MAF <0.01, and tested independently.

| Supplementary Table 6: Nominally significant results from gene-based association |  |  |  |  |  |  |
| --- | --- | --- | --- | --- | --- | --- |
| LOCUS | TYPE | LOCATION | CTRL_FREQ | CASE_FREQ | P<sub>locus</sub> | P<sub>corr</sub> |
| **NRXN1** | DEL | chr2:50145642-51259674 | 12 | 1 | 6.2e-5 | 6.7e-4 |
| **CALY** | DEL | chr10:135138927-135150475 | 4 | 0 | 0.019 | 0.50 |
| **PRAP1** | DEL | chr10:135160843-135166187 | 4 | 0 | 0.019 | 0.50 |
| **ZNF511** | DEL | chr19:53935226-53947925 | 4 | 0 | 0.019 | 0.50 |
| **CNTN6** | DUP | chr3:1134341-1445292 | 12 | 2 | 2.5e-4 | 8.3e-3 |
| **CNTN4** | DUP | chr3:2140549-3099645 | 8 | 3 | 0.018 | 0.93 |
| **SHCBP1** | DUP | chr16:46614467-46655311 | 4 | 0 | 0.019 | 0.90 |
| **VPS35** | DUP | chr16:46693588-46723144 | 4 | 0 | 0.019 | 0.90 |
| **ORC6** | DUP | chr16:46723557-46732306 | 4 | 0 | 0.019 | 0.90 |
| **IGSF11** | DUP | chr3:118619478-118864898 | 18 | 15 | 0.032 | 1.0 |
| **SPSB1** | DUP | chr1:9352940-9429590 | 25 | 25 | 0.044 | 1.0 |
| **RPL38** | DUP | chr17:72199721-72206794 | 12 | 9 | 0.050 | 1.0 |
| **TTYH2** | DUP | chr17:72209653-72258155 | 12 | 9 | 0.050 | 1.0 |
| **DNAI2** | DUP | chr17:72270386-72311023 | 12 | 9 | 0.050 | 1.0 |
| **KIF19** | DUP | chr17:72322349-72351959 | 12 | 9 | 0.050 | 1.0 |
| **BTBD17** | DUP | chr17:72352555-72358085 | 12 | 9 | 0.050 | 1.0 |
| **GPR142** | DUP | chr17:72363546-72368761 | 12 | 9 | 0.050 | 1.0 |
| **GPRC5C** | DUP | chr17:72420990-72447792 | 12 | 9 | 0.050 | 1.0 |
| **CD300A** | DUP | chr17:72462555-72480933 | 12 | 9 | 0.050 | 1.0 |
| **CD300LB** | DUP | chr17:72517313-72527613 | 12 | 9 | 0.050 | 1.0 |
| **CD300C** | DUP | chr17:72537247-72542282 | 12 | 9 | 0.050 | 1.0 |
| **CD300LD** | DUP | chr17:72575504-72588422 | 12 | 9 | 0.050 | 1.0 |
| **C17orf77** | DUP | chr17:72580818-72590348 | 12 | 9 | 0.050 | 1.0 |
| **CD300E** | DUP | chr17:72606026-72619897 | 12 | 9 | 0.050 | 1.0 |

**Supplementary Figure 8.**
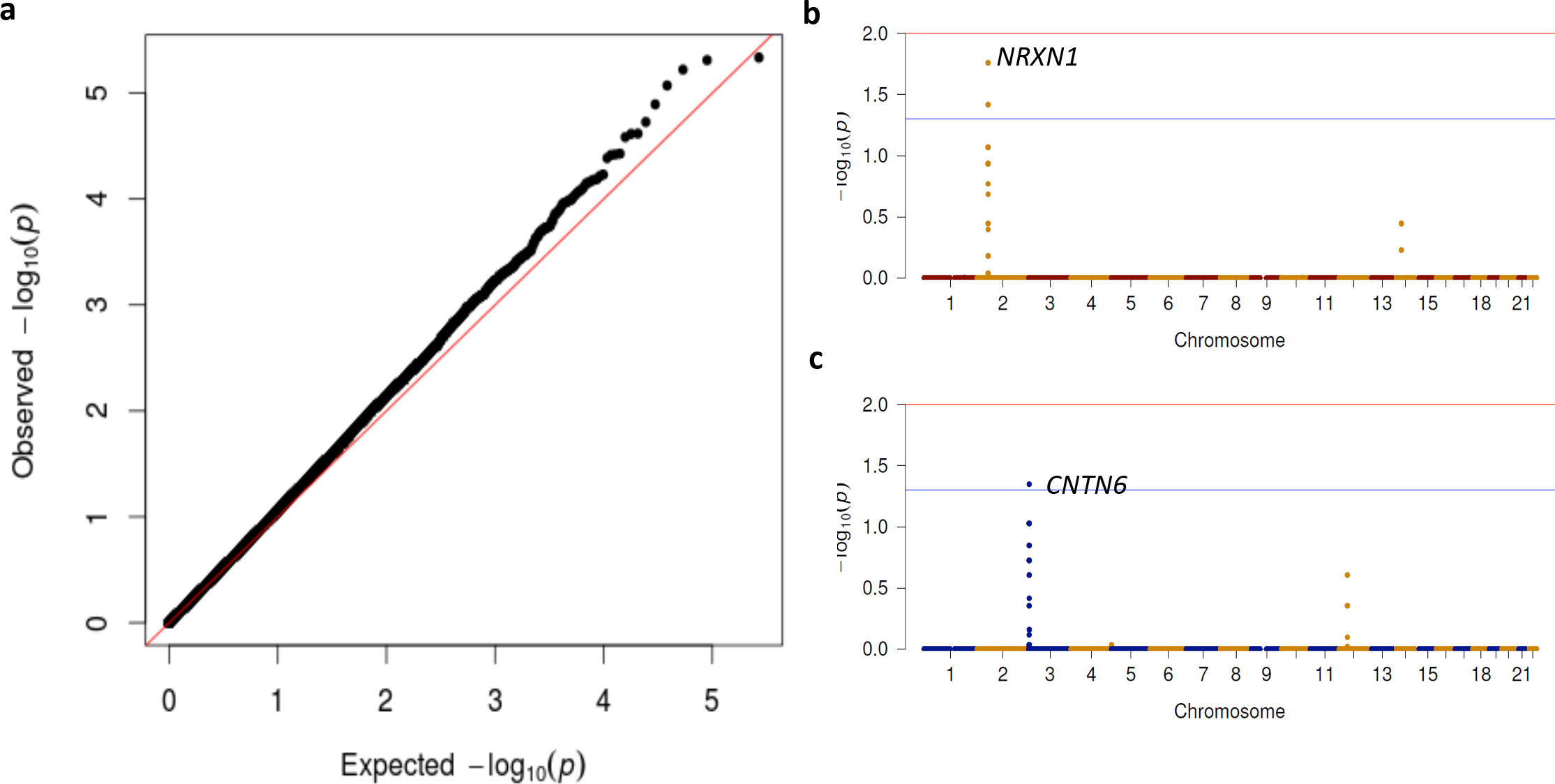
Examination for population-specific effects. To verify the robustness of our results to population stratification, we pair-matched each case subject with exactly one case such that the global difference between all pairs is minimized using SpectralGEM. We also excluded all outlying pairs based on the genetic distance between them (>90th percentile). (a) The SNP-based λgc of the resultant dataset (1996 cases and 1996 controls) was an acceptable 1.082. Manhattan plots of segmental association results demonstrate that (b) deletions in *NRXN_1_* and (c) duplications in *CNTN6* are significant with an a < 0.05 (blue line). Deletions and duplications were analyzed separately. The −log10 (p-value) displayed is empirically corrected for FWER genome-wide using the max(T) method with 1,000,000 permutations.

**Supplementary Table 7.** Prevalence of known, recurrent, neuropsychiatric CNVs in TS. Known CNVs were considered present in a sample if it carried a variant of the same CNV type (deletion/duplication) and overlapped at least 50% the length of the known recurrent CNV, except for the single-gene locus containing *VIPR_2_*, where the associated signal has been shown to derive from non-recurrent events^37^. For this locus, any CNV > 50kb overlapping any portion of the gene was counted. P-values were determined with 100,000 permutations using the max(T) method to control for the family-wise error rate. P-values corrected empirically for multiple testing are shown in parentheses, with two-sided p-values presented only when there is enrichment in controls.

| Supplementary Table 7: Prevalence of known, recurrent, pathogenic CNVs in TS |  |  |  |  |  |  |
| --- | --- | --- | --- | --- | --- | --- |
| LOCUS | LOCATION (hg19) | CNV TYPE | CASE FREQ | CTRL FREQ | EMP-P (1-sided) | EMP-P (2-sided) |
| **1q21.1** | 1:146089254-147859944 | Loss | 1 | 1 | 0.62(1) |  |
| **3q29** | 3:195766737-197216349 | Loss | 0 | 0 | 1(1) |  |
| **7q36.3 (VIPR2)** | 7:157860945-159119486 | Loss | 0 | 0 | 1(1) |  |
| **10q11.22-23** | 10:46508694-51912781 | Loss | 0 | 0 | 1(1) |  |
| **15q11.2** | 15:22750305-23272733 | Loss | 10 | 8 | 0.1(0.59) |  |
| **15q13.1** | 15:28973396-30556183 | Loss | 0 | 0 | 1(1) |  |
| **15q13.3** | 15:30936285-32620127 | Loss | 0 | 1 | 1(1) |  |
| **16p13.11** | 16:15125441-16291983 | Loss | 1 | 1 | 0.62(1) |  |
| **16p12.1** | 16:21947230-22423698 | Loss | 1 | 1 | 0.62(1) |  |
| **16p11.2 distal** | 16:28814098-29043450 | Loss | 0 | 0 | 1(1) |  |
| **16p11.2** | 16:29595483-30192561 | Loss | 2 | 2 | 0.5(0.98) |  |
| **17p12** | 17:14101029-15471179 | Loss | 1 | 0 | 0.39(0.94) |  |
| **17q12** | 17:34815551-36249430 | Loss | 0 | 1 | 1(1) | 1(1) |
| **22q11.21 short** | 22:18905281-21488855 | Loss | 0 | 0 | 1(1) |  |
| **22q11.21 long** | 22:20733495-21465780 | Loss | 1 | 0 | 0.39(0.94) |  |
| **1q21.1** | 1:146089254-147859944 | Gain | 2 | 3 | 0.63(1) | 1(1) |
| **2p25.3** | 2:1754610-2225144 | Gain | 0 | 0 | 1(1) |  |
| **7q36.3 (VIPR2)** | 7:158726462-158947294 | Gain | 0 | 1 | 1(1) | 1(1) |
| **10q11.22-23** | 10:47543322-51912781 | Gain | 0 | 0 | 1(1) |  |
| **13q31.3** | 13:94358193-94419894 | Gain | 0 | 0 | 1(1) |  |
| **15q11.2-13.1** | 15:22770994-28535266 | Gain | 2 | 0 | 0.15(0.35) |  |
| **15q13.1** | 15:29562640-30689724 | Gain | 0 | 0 | 1(1) |  |
| **16p13.11** | 16:14989844-16291983 | Gain | 1 | 6 | 1(1) | 0.26(0.73) |
| **16p11.2** | 16:29624247-30198151 | Gain | 0 | 1 | 1(1) | 1(1) |
| **17q12** | 17:34815551-36249430 | Gain | 2 | 2 | 0.5(0.94) |  |
| **22q11.21 short** | 22:18905281-21488855 | Gain | 3 | 4 | 0.54(0.99) |  |
| **22q11.21 long** | 22:20718116-21465780 | Gain | 0 | 0 | 1(1) |  |

**Supplementary Table 8.** Clinical phenotypes of *NRXN_1_* and *CNTN6* CNV carriers. Clinical phenotypes for all CNV carriers of the two significant TS loci detected in this study (gene-based association test): Deletions at *NRXN_1_* and duplications at *CNTN6*, including common comorbid disorders for TS, attention deficit disorder (ADHD) and obsessive compulsive disorder (OCD) as well as whether or not the individual was flagged for an atypical phenotype (FLAGGED, and described in NOTES).

| SAMPLE_ID | GENE | CHR | START | END | CNV_TYPE | LENGTH_KB | CNV_EFFECT | OCD | ADHD | FLAGGED | NOTES |
| --- | --- | --- | --- | --- | --- | --- | --- | --- | --- | --- | --- |
| **TS1_0630** | NRXN1 | 2 | 50821559 | 51021488 | DEL | 199.9 | CODING | 1 | 1 | Y | Unspecified Developmental Delay (ICD-9: 315.9) |
| **TS1_0180** | NRXN1 | 2 | 50930181 | 51272375 | DEL | 342.2 | CODING | 1 | 1 | Y | Asperger Syndrome |
| **TS1_0446** | NRXN1 | 2 | 50945471 | 51770480 | DEL | 825 | CODING | 2 | 2 | N |  |
| **TS1_0105** | NRXN1 | 2 | 51002606 | 51316822 | DEL | 314.2 | CODING | 1 | 2 | N |  |
| **TS2_1256** | NRXN1 | 2 | 51028662 | 51458570 | DEL | 429.9 | CODING | 1 | 2 | Y | Other developmental speech or language disorder (ICD-9: 315.39) |
| **TS2_0026** | NRXN1 | 2 | 51041472 | 51483528 | DEL | 442.1 | CODING | 1 | 1 | N |  |
| **TS2_0924** | NRXN1 | 2 | 51041603 | 51528298 | DEL | 486.7 | CODING | 1 | 2 | N |  |
| **TS2_0750** | NRXN1 | 2 | 51058745 | 51252137 | DEL | 193.4 | CODING | 2 | 2 | Y | Asperger Syndrome |
| **TS2_1238** | NRXN1 | 2 | 51077569 | 51458570 | DEL | 381 | CODING | 2 | 1 | Y | Paranoid personality disorder |
| **TS1_0573** | NRXN1 | 2 | 51079482 | 51357902 | DEL | 278.4 | CODING | NA | NA | NA |  |
| **TS1_0776** | NRXN1 | 2 | 51101583 | 51308895 | DEL | 207.3 | CODING | 1 | 2 | N | Brother with Asperger Syndrome |
| **TS1_0698** | NRXN1 | 2 | 51123048 | 51286169 | DEL | 163.1 | CODING | 2 | 2 | N |  |
| **TS2_1805** | CNTN6 | 3 | 565961 | 1350458 | DUP | 784.5 | CODING | 2 | NA | N |  |
| **TS2_1405** | CNTN6 | 3 | 668832 | 1143424 | DUP | 474.6 | 5' UTR | 2 | 2 | N |  |
| **TS2_1624** | CNTN6 | 3 | 707257 | 1781739 | DUP | 1074 | CODING | 1 | 1 | N |  |
| **TS2_1525** | CNTN6 | 3 | 857325 | 1427769 | DUP | 570.4 | CODING | 2 | 2 | N |  |
| **TS2_1568** | CNTN6 | 3 | 864513 | 1425997 | DUP | 561.5 | CODING | 2 | 2 | N |  |
| **TS2_1545** | CNTN6 | 3 | 864513 | 1427769 | DUP | 563.3 | CODING | 1 | 2 | N |  |
| **TS2_1320** | CNTN6 | 3 | 946290 | 1276092 | DUP | 329.8 | CODING | 2 | 1 | N |  |
| **TS1_0618** | CNTN6 | 3 | 1125605 | 1315900 | DUP | 190.3 | CODING | 1 | 1 | N |  |
| **TS2_1156** | CNTN6 | 3 | 1218279 | 2170519 | DUP | 952.2 | CODING | 1 | 2 | N |  |
| **TS1_0558** | CNTN6 | 3 | 1218279 | 2170519 | DUP | 952.2 | CODING | 1 | 2 | N |  |
| **TS2_0827** | CNTN6 | 3 | 1226953 | 2170519 | DUP | 943.6 | CODING | 1 | 1 | N |  |
| **TS2_0452** | CNTN6 | 3 | 1260932 | 1556680 | DUP | 295.7 | CODING | 1 | 1 | N |  |

**Supplementary Figure 9.**
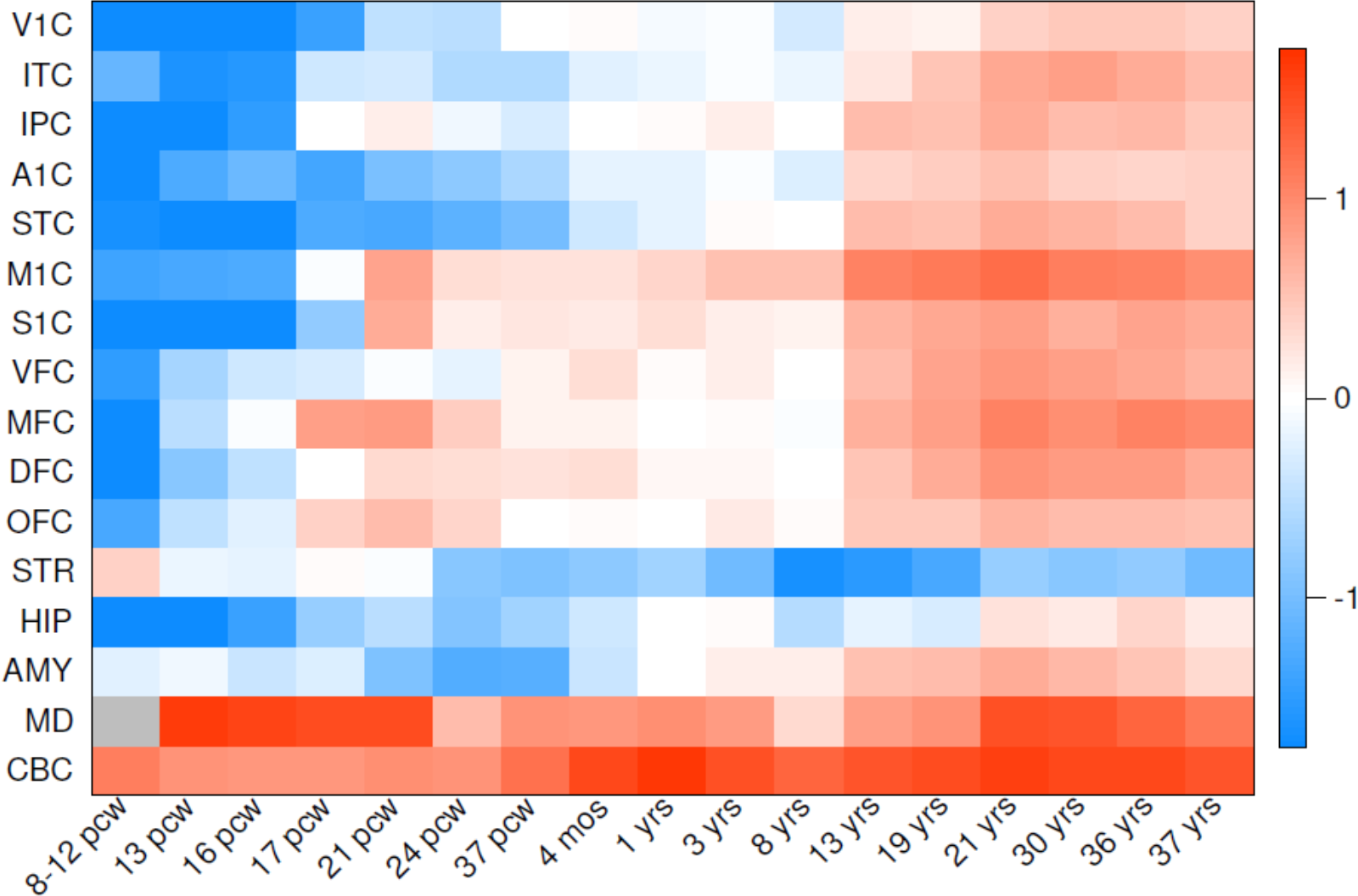
Spatio-temporal expression of *CNTN6* in human brain. Analysis of RNASeq data from Brainspan (www.brainspan.org) indicates that *CNTN6* is most highly expressed postnatally in the cerebellum (CBC) followed by the thalamus (MD). Heatmap represents median expression values for all available samples at each tissue normalized across each developmental timepoint. Regions: primary visual cortex (V1C), inferolateral temporal cortex (ITC), posteroventral parietal cortex (IPC), primary auditory cortex (A1C), posterior superior temporal cortex (STC), primary motor cortex (M1C), primary somatosensory cortex (S1C), ventrolateral prefrontal cortex (VFC), anterior cingulate cortex (MFC), dorsolateral prefrontal cortex (DFC), orbital frontal cortex (OFC), striatum (STR), hippocampus (HIP), amygdaloid complex (AMY), mediodorsal nucleus of thalamus (MD), cerebellar cortex (CBC). Temporal abbreviations: post-conception weeks (pcw), months (mos), years (yrs).

**Supplementary Table 9.**
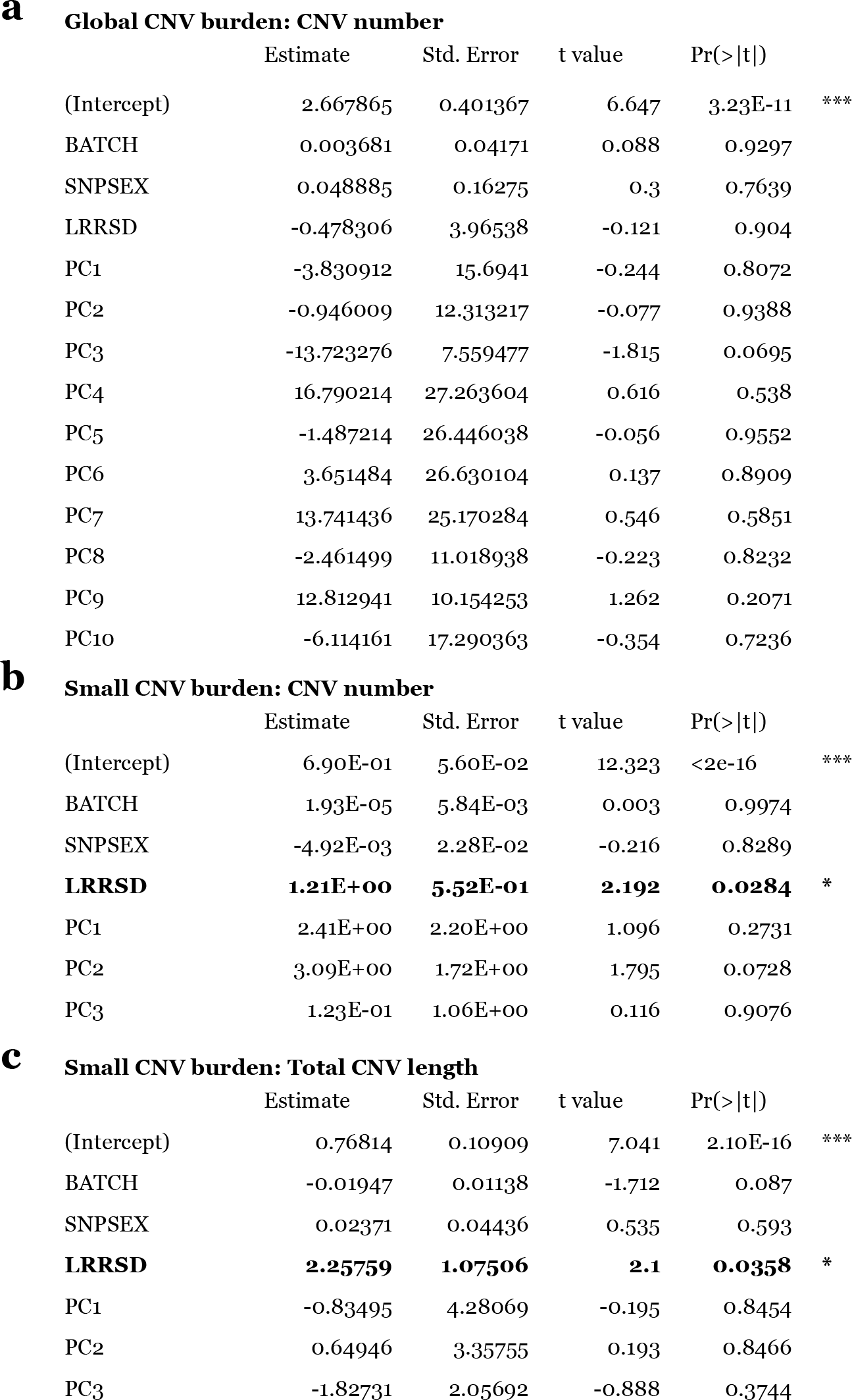
Regression models of global CNV burden. (a) Results from a linear regression model of global burden with total CNV count as the dependent variable, and genotyping batch, subject sex, LogR-Ratio standard deviation (LRRSD, a general metric for CNV assay quality), and the top 10 ancestry PCs as dependent variables. None of the included covariates were significant predictors of global CNV burden in terms of CNV count or total CNV size (Supplementary Text). However, when restricted to small CNV events (<100kb), LRRSD was a significant predictor of CNV burden in terms of (b) total CNV count and (c) total CNV length; therefore LRRSD was included as a covariate in our subsequent analysis.

| **a Global CNV burden: CNV number** |  |  |  |  |  |
| --- | --- | --- | --- | --- | --- |
|  | Estimate | Std. Error | t value | Pr(>\|t\|) |  |
| (Intercept) | 2.667865 | 0.401367 | 6.647 | 3.23E-11 | *** |
| BATCH | 0.003681 | 0.04171 | 0.088 | 0.9297 |  |
| SNPSEX | 0.048885 | 0.16275 | 0.3 | 0.7639 |  |
| LRRSD | -0.478306 | 3.96538 | -0.121 | 0.904 |  |
| PC1 | -3.830912 | 15.6941 | -0.244 | 0.8072 |  |
| PC2 | -0.946009 | 12.313217 | -0.077 | 0.9388 |  |
| PC3 | -13.723276 | 7.559477 | -1.815 | 0.0695 |  |
| PC4 | 16.790214 | 27.263604 | 0.616 | 0.538 |  |
| PC5 | -1.487214 | 26.446038 | -0.056 | 0.9552 |  |
| PC6 | 3.651484 | 26.630104 | 0.137 | 0.8909 |  |
| PC7 | 13.741436 | 25.170284 | 0.546 | 0.5851 |  |
| PC8 | -2.461499 | 11.018938 | -0.223 | 0.8232 |  |
| PC9 | 12.812941 | 10.154253 | 1.262 | 0.2071 |  |
| PC10 | -6.114161 | 17.290363 | -0.354 | 0.7236 |  |
| **b Small CNV burden: CNV number** |  |  |  |  |  |
|  | Estimate | Std. Error | t value | Pr(>\|t\|) |  |
| (Intercept) | 6.90E-01 | 5.60E-02 | 12.323 | <2e-16 | *** |
| BATCH | 1.93E-05 | 5.84E-03 | 0.003 | 0.9974 |  |
| SNPSEX | -4.92E-03 | 2.28E-02 | -0.216 | 0.8289 |  |
| **LRRSD** | **1.21E+00** | **5.52E-01** | **2.192** | **0.0284** | * |
| PC1 | 2.41E+00 | 2.20E+00 | 1.096 | 0.2731 |  |
| PC2 | 3.09E+00 | 1.72E+00 | 1.795 | 0.0728 |  |
| PC3 | 1.23E-01 | 1.06E+00 | 0.116 | 0.9076 |  |
| **c Small CNV burden: Total CNV length** |  |  |  |  |  |
|  | Estimate | Std. Error | t value | Pr(>\|t\|) |  |
| (Intercept) | 0.76814 | 0.10909 | 7.041 | 2.10E-16 | *** |
| BATCH | -0.01947 | 0.01138 | -1.712 | 0.087 |  |
| SNPSEX | 0.02371 | 0.04436 | 0.535 | 0.593 |  |
| **LRRSD** | **2.25759** | **1.07506** | **2.1** | **0.0358** | * |
| PC1 | -0.83495 | 4.28069 | -0.195 | 0.8454 |  |
| PC2 | 0.64946 | 3.35755 | 0.193 | 0.8466 |  |
| PC3 | -1.82731 | 2.05692 | -0.888 | 0.3744 |  |

